# Distinct architectural requirements for the *parS* centromeric sequence of the pSM19035 plasmid partition machinery

**DOI:** 10.1101/2022.05.29.493902

**Authors:** Andrea Volante, Juan C. Alonso, Kiyoshi Mizuuchi

**Affiliations:** Laboratory of Molecular Biology, National Institute of Diabetes and Digestive and Kidney Diseases, National Institutes of Health, USA; Departamento de Biotecnología Microbiana, Centro Nacional de Biotecnología, Consejo Superior de Investigaciones, Científicas, 28049 Madrid, Spain; Merck, 39 Rte Industrielle de la Hardt, Molsheim 67120, France

**Keywords:** Plasmid partition, ParABS system, Chromosome segregation

## Abstract

Three-component ParABS partition systems ensure stable inheritance of many bacterial chromosomes and low-copy-number plasmids. ParA localizes to the nucleoid through its ATP- dependent non-specific DNA binding activity, whereas centromere-like *parS*-DNA and ParB form partition complexes that activate ParA-ATPase to drive the system dynamics. The essential *parS* sequence arrangements vary among ParABS systems, reflecting the architectural diversity of their partition complexes. Here, we focus on the pSM19035 plasmid partition system that uses a ParB_pSM_ of the ribbon-helix-helix (RHH) family. We show that *parS_pSM_* with four or more contiguous ParB_pSM_-binding sequence repeats is required to assemble a stable ParA_pSM_-ParB_pSM_ complex and efficiently activate the ParA_pSM_-ATPase, stimulating complex disassembly. Disruption of the contiguity of the *parS_pSM_* sequence array destabilizes the ParA_pSM_-ParB_pSM_ complex and prevents efficient ATPase activation. Our findings reveal the unique architecture of the pSM19035 partition complex and how it interacts with nucleoid-bound ParA_pSM_-ATP.

## Introduction

Faithful chromosome segregation is essential for the proliferation of bacterial cells, and low-copy-number plasmids also need a robust partition mechanism for their stable inheritance. However, prokaryotes do not possess the mitotic machinery of eukaryotes: instead, alternative active DNA partition systems have evolved, among which ParABS systems (also called class I partition systems) are the most widespread. Basic ParABS systems consist of three components, a partition ATPase (ParA), a “centromere” binding protein (ParB), and a *cis*-acting centromere-like DNA site (*parS*).

ATP-activated ParA dimers bind nonspecific DNA (nsDNA) and localize to the nucleoid *in vivo* (Ebersbach and Gerdes 2001, Ebersbach and Gerdes 2004, Pratto et al. 2008, Ringgaard et al. 2009). The *parS* sites, often composed of multiple tandem repeats of binding consensus sequences for ParBs, demark the DNA-cargos that are translocated and positioned into the two halves of the cell before cell division by recruiting ParB molecules to assemble partition complexes (PCs). ParB proteins fall into two structurally unrelated groups, dimeric helix-turn-helix (HTH), and dimeric ribbon-helix-helix (RHH) DNA binding proteins. HTH-ParBs have been shown to bind not only site-specifically to their cognate *parS* sequences but also to spread many kilobase pairs into the DNA neighboring the *parS* sites (Murray et al. 2006, Graham et al. 2014, Soh et al. 2019, Rodionov et al. 1999, Lynch and Wang 1995, Breier and Grossman 2007, Jalal et al. 2020, Osorio-Valeriano et al. 2019, Sanchez et al. 2015). Therefore, they form large PCs containing many ParB molecules bound to condensed DNA around *parS*. In contrast, RHH-ParBs are not known to spread into non-specific DNA flanking *parS*, though like HTH-ParBs, they interact with their cognate ParA proteins *via* their N-terminus (Radnedge et al. 1998, Figge et al. 2003, Barilla et al. 2007). Many RHH-ParB proteins also control the expression of the proteins involved in the partition system and plasmid copy number control by binding *parS* sites, which overlaps promotors of their genes (de la Hoz et al. 2000).

ParA-ParB interaction leads to activation of the ParA-ATPase, which is most efficient in the presence of nsDNA and *parS* DNA (Ah-Seng et al. 2009; Chu et al. 2019; Pratto et al. 2008; Taylor et al., 2021) and leads to dissociation of ParA from nsDNA. Interaction dynamics between the nucleoid-bound ParA and ParB in the PC prior to ATP hydrolysis and ParA dissociation determine the dynamics of the PC relative to the nucleoid. The common results of most systems *in vivo* appear to be the establishment of equidistant distribution of two or more PCs along the nucleoid so that at cell division, each daughter cell inherits at least one copy of the plasmid DNA (Sengupta et al. 2010, Ringgaard et al. 2009, Lioy et al. 2015, McLeod et al. 2017).

With biochemical findings and observations from live-cell imaging approaches accumulating in the field, combined with experiments using reconstituted cell-free reaction systems, a diffusion- ratchet mechanism of the ParABS partition was proposed. Here, the driving force for the DNA- cargo motion is generated by a propagating nucleoid-bound ParA distribution gradient (Vecchiarelli et al. 2010, Hwang et al. 2013, Vecchiarelli et al. 2013, Vecchiarelli et al. 2014). Additional models related to the diffusion-ratchet mechanism have also been proposed based on high-resolution imaging observations (Lim et al. 2014, Le Gall et al. 2016). While findings supporting diffusion-ratchet-type models for ParABS systems accumulate, many molecular details required to put the model on quantitatively solid ground are lacking. ParB “spreading”, a unique feature of PCs involving HTH-ParB proteins, loads many ParB molecules onto a PC to facilitate partitioning. However, it is unclear if a similar mechanism could explain the systems with RHH-ParBs. Nevertheless, live-cell imaging studies of systems that use RHH-ParB proteins showed an oscillating dynamic ParA distribution pattern on the nucleoid and PC chasing the receding tail of the ParA distribution (Ringgaard et al. 2009, Lioy et al. 2015, McLeod et al. 2017) similar to observations with the F-plasmid partition system and other related systems involving HTH-ParB proteins (Hatano and Niki 2010, Schofield et al. 2010) for which there is accumulating evidence supporting the diffusion-ratchet mechanism. Therefore, we suspect systems with RHH-ParBs also operate *via* a diffusion-ratchet-type mechanism. A better understanding of how the PCs are organized for this group and a quantitative understanding of the interaction dynamics of these PCs with nucleoid-bound ParA molecules are critical for advancing our mechanistic understanding of these systems.

In this study, we focused on the pSM19035 partition system of *Streptococcus pyogenes*. This plasmid harbors a ParABS system composed of ParA_pSM_ (also called Delta), an RHH ParB_pSM_ (also called Omega), and six *parS_pSM_* sites, each comprising 7-10 consecutive non-palindromic 7- bp-long sequence repeats (5’-WATCACW-3’, symbolized by →) that overlap the promoter regions of *copS*, δ (coding ParA_pSM_) and ω (coding ParB_pSM_) genes. Each ParB_pSM_ dimer binds one copy of the 7-bp *parS_pSM_* consensus sequence. However, the affinity for a single repeat is low, while two dimers bind with high affinity to two direct (→→) or inverted (→←) repeats forming dimers of dimers (de la Hoz et al. 2004, Weihofen et al. 2006, Welfle et al. 2005). Within this ParB_pSM_-*parS_pSM_* complex, DNA does not show significant curvature (Weihofen et al. 2006) and full-size *parS_pSM_*-ParB_pSM_ complexes also appear to maintain a straight DNA configuration around which ParB_pSM_ dimers are predicted to form short left-handed spiral (Pratto et al. 2008, Pratto et al. 2009).

ParA_pSM_, unlike most other ParAs, forms dimers in solution in the absence of ATP (Pratto et al. 2008). Like other ParAs, it also undergoes a conformational transition upon binding ATP that increases its affinity for nsDNA (Soberon et al. 2011, Pratto et al. 2008). In the ATP-bound form, ParA_pSM_ has been shown to bind nsDNA forming short clustered patches containing several ParA_pSM_ dimers at random location, instead of individual dimers independently distributed on the nsDNA (Pratto et al. 2009). In the presence of *parS_pSM_* DNA and ParB_pSM_, several *parS_pSM_*-ParB_pSM_ mini-filaments and ParA_pSM_-nsDNA patches appeared to bind together to form large protein-DNA complexes bridging multiple DNA molecules (Pratto et al. 2008, Pratto et al. 2009, Soberon et al. 2011, Lioy et al. 2015). Interactions among these components fueled by ATP hydrolysis are thought to drive dynamic oscillations of the nucleoid-bound ParA_pSM_ *in vivo*, which resembles those observed for the TP228 ParABS system (Lioy et al. 2015, McLeod et al. 2017). However, how the observed inter-molecular interactions coordinate the *in vivo* system dynamics resulting in robust plasmid partitioning remains a mystery. To approach this puzzle, here we studied functional requirements of the *parS_pSM_* sequence-structure necessary for ParB_pSM_-mediated activation of the ParA_pSM_ ATPase. We found a minimum of four contiguous repeats of the *parS_pSM_* heptad consensus sequence without a gap is necessary for full activity. Kinetics of the nsDNA-bound *parS_pSM_*-ParB_pSM_-ParA_pSM_ complex formation and disassembly indicated the presence of a complex multistep process involved in ATPase activation.

## Results

### ParA_pSM_ ATPase is synergistically activated by nsDNA, ParB_pSM_ and *parS_pSM_*-DNA

To define the requirements for stimulation of the ParA_pSM_ ATPase activity by ParB_pSM_, the steady- state ParA_pSM_ ATP turnover rate was measured with varying concentrations of ParB_pSM_ in the presence or absence of different duplex DNA cofactors. The turnover rate of ParA_pSM_ ATPase alone is very low at 37°C (0.9 ± 0.1 ATP/ParA-dimer/h, N=3; Fig. 1A; no DNA). In the presence of a saturating concentration of double stranded DNA (40 μg/ml pBR322 plasmid DNA plus 23 to 38 μg/ml double stranded oligonucleotide with or without *parS_pSM_* sequence), to which ATP- ParA_pSM_ dimers can bind to support ATPase activation by ParB_pSM_ (see below), no significant rate change was observed (1.0 ± 0.1 h_-1_, N=46, Fig. 1A, pool of all measurements at [ParB_pSM_] = 0). Next, effects of the *parS_pSM_*-DNA in addition to 40 μg/ml nsDNA and ParB_pSM_ were examined. ParA_pSM_ ATP hydrolysis was stimulated up to ∼20-fold (*k_cat_* = 20.5 ± 2.9 h_-1_, N=7) in the presence of ParB_pSM_ and oligonucleotide duplex DNA containing one of the native arrangements containing seven *parS_pSM_* heptad-sequence-repeats (7R-*parS_pSM_*, →→←→→←←) (Fig. 1A). The stimulation approached saturation around 2 μM ParB_pSM_ at varying ParA_pSM_ concentrations (Fig. 1-figure supplement 1A). In contrast, when the *parS_pSM_* DNA fragment was replaced with one having a scrambled sequence, ParB_pSM_ stimulated ParA_pSM_ ATPase activity only to 2.4 ± 1.1 h_-1_ even in the presence of 8 μM ParB_pSM_ that should have allowed non-*parS_pSM_* DNA binding (N=3*) (Fig. 1A, scram). Similarly, a low level of ParA_pSM_ ATPase stimulation was observed with ParB_pSM1-27_ peptide, which lacked the DNA binding and dimerization domains (Fig. 1-figure supplement 1B). These results demonstrated that efficient ParA_pSM_-ATPase stimulation by ParB_pSM_ requires specific *parS_pSM_* interactions. Even the low *parS_pSM_*-independent ATPase stimulation was not detected in the absence of DNA (Fig. 1A, no DNA). ParB_pSM1-27_ peptide was also unable to stimulate the ATPase without DNA (Fig. 1-figure supplement 1B). We conclude ParA_pSM_-nsDNA binding is required for the ATPase activation by ParB_pSM_, as reported for ParA_F_ ATPase activation by ParB_F_ (Taylor, et al 2021). Consistently, in the absence of nsDNA, excess ParB_pSM_ relative to the concentration of 7R-*parS_pSM_* in the reaction inhibited ATPase stimulation, presumably competing with ParA_pSM_ for *parS_pSM_* DNA binding (Fig. 1-figure supplement 1C).

**Figure 1.**
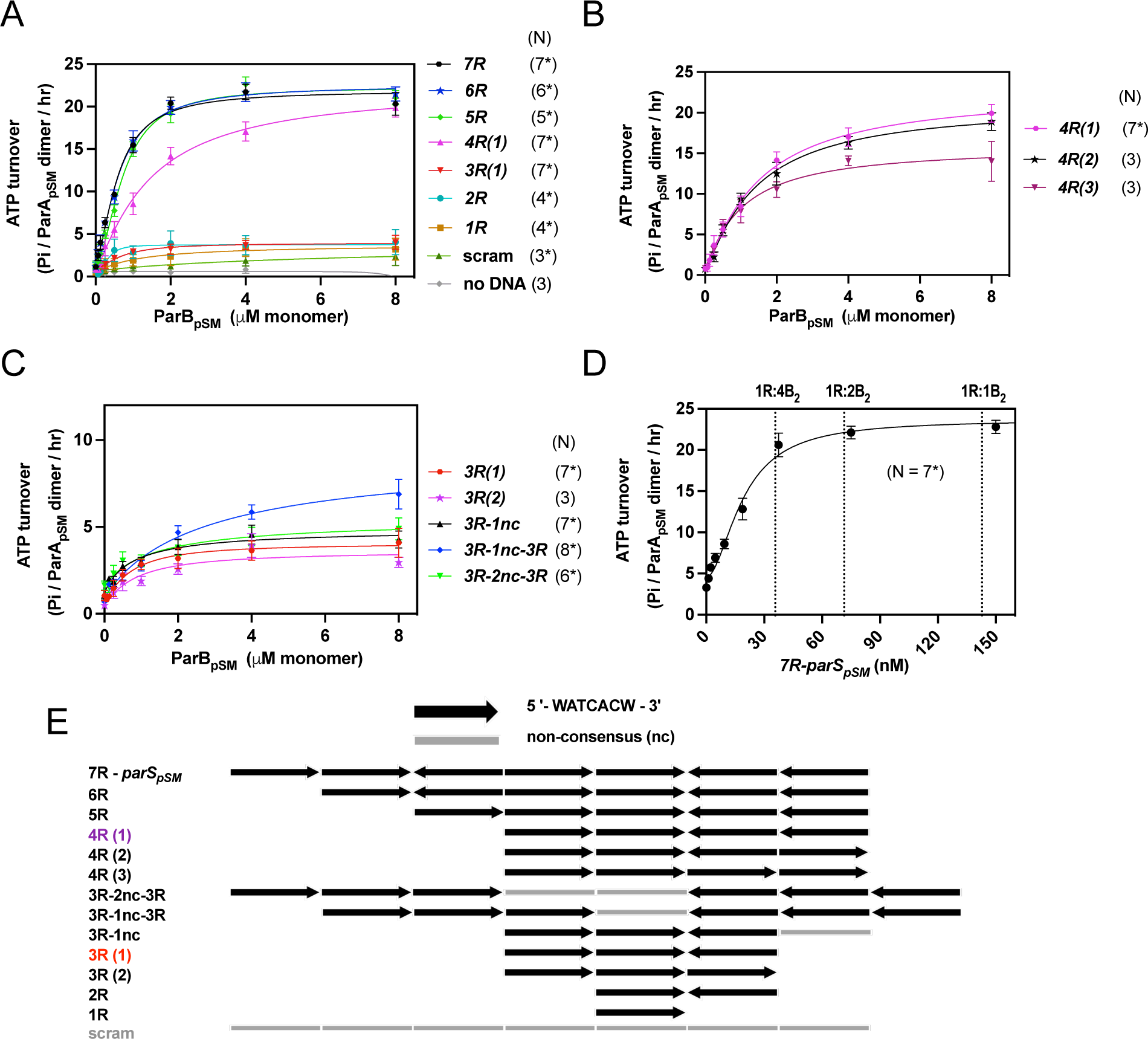
ParA_pSM_ ATPase activation by ParB_pSM_, *parS_pSM_* and nsDNA. (A) Efficient activation of ParA**_pSM_** ATPase by ParB**_pSM_** exhibits critical dependency on *parS_pSM_* heptad sequence repeat number. The ATPase reactions contained ParA_pSM_ (2μM), pBR322 DNA (60 μM in bp, unless noted otherwise), ParB_pSM_ (at the concentration indicated) and *parS_pSM_* duplex substrates (4.4 μM of the 7-bp consensus sequence repeats) or equal amount of a scrambled sequence duplex (scram). (B) Comparison of *parS_pSM_* containing 4 contiguous repeats with different heptad orientation arrangements. (C) *parS_pSM_* containing different heptad arrangements of 3 contiguous repeats and two contiguous triple heptad repeats with a gap fails to fully activate the ParA_pSM_ ATPase. (D) *7R-parS_pSM_* concentration dependence of ParA_pSM_ ATPase activity. Reaction mixtures contained ParA_pSM_ (2μM), ParB_pSM_ (2μM), pBR322 DNA (60 μM in bp), and increasing concentration of *7R-parS_pSM_* duplex (duplex fragment concentration shown, the ratio of *parS_pSM_* heptad repeat sequence to ParB_pSM_ dimers are indicated on top). (E) The numbers and arrangement of the heptad repeats of the *parS_pSM_* fragments used in this study (also see Fig. S1). Data points represent means and standard errors of mean (SEM) of N repeated experiments (N* represents repeats for majority of data points, see Source Data File for details). Curves were fitted after subtraction of the background in the absence of ParA_pSM_ to an equation v-v_0_=(v_max_[B]_n_)/(K_An_+[B]_n_). The maximum turnover rates (v_max_) cited in the text represent mean ± 95% confidence intervals (symmetrized to the larger estimated errors from the mean for simplicity). Source Data 1 Figure supplement 1: ParA_pSM_-ATPase stimulation: by ParB_pSM_ at different ParA_pSM_ concentrations, by ParB_pSM1-27_, and by ParB_pSM_-7R-*parS_pSM_* in the absence of nsDNA. Figure supplement 2: DNA duplex substrates. Figure supplement 3: Binding affinity of ParB_pSM_ to 3R- and 4R-*parS_pSM_*.

### Efficient ParA_pSM_-ATPase stimulation requires ParB_pSM_ bound to *parS_pSM_*-DNA with at least 4 contiguous heptad-sequence-repeats

We asked if entire 7R-*parS_pSM_* is required for the full stimulation of ParA_pSM_ ATPase by ParB_pSM_. A series of deletions of *parS_pSM_* heptad sequence repeats were made and their ATPase stimulation activities were tested (Fig. 1E). Efficient ATPase stimulation was observed when 6R (→←→→←←), 5R (→→→←←), or 4R (→→←←) was added along ParB_pSM_ (Fig. 1A). No significant difference was observed among 5R, 6R or 7R; both half-saturation concentrations and the apparent *k_cat_* were comparable (Fig. 1A). ParB_pSM_-4R-*parS_pSM_* also induced similar rates of ParA_pSM_ ATP hydrolysis, but a significantly higher concentration was required for full stimulation (Fig. 1A). Different heptad orientation arrangements of 4R-*parS_pSM_* (→→←→ and →→→→) stimulated ParA_pSM_ ATPase to a similar extent (Fig. 1B). In contrast, 3R- (→→←), 2R- (→←), and 1R- (→) *parS_pSM_* were poor cofactors for ParB_pSM_-dependent ATPase stimulation (apparent *k_cat_* = 3.2-3.6 ± 1.3-2.1 h_-1_, N=4-7*, Fig. 1A), not significantly different from the scrambled sequence DNA. 3R-*parS_pSM_* fragments with different repeat arrangements behaved similarly to each other (Fig. 1C). The ATPase stimulation by *parS_pSM_* DNA with 1 to 3 *parS_pSM_* repeats also appeared to saturate at around 2 μM ParB_pSM_, indicating that the affinity of the nsDNA-bound ParA_pSM_ to ParB_pSM_ in the presence of truncated *parS_pSM_* was not limiting above ∼2 μM ParB_pSM_. Previously it has been shown that ParB_pSM_ bound 4R or 3R DNA fragments with a similar affinity that was >50-fold higher than nsDNA (de la Hoz et al. 2004). We confirmed that ParB_pSM_ binding was strong for 4R and 3R duplex DNA (*K*_D_ ∼17 nM) and weak for nsDNA (*K*_D_ ∼1μM, Fig. 1- figure supplement 3). Therefore, the affinity of ParB_pSM_ for the 3R-*parS_pSM_* DNA is not limiting the ParA_pSM_ ATPase stimulation. In the above experiment, the length of *parS_pSM_* DNA fragments decreased as the number of *parS_pSM_* repeats decreased (Figs. 1E, Fig. 1-figure supplement 2). We tested longer 3R-*parS_pSM_* DNA fragments containing an additional non-*parS_pSM_* heptamer sequence (non-consensus, nc) (3R-1nc) and confirmed that *parS_pSM_*-DNA fragment size was not a significant factor for the ATPase stimulation efficiency (Fig. 1C).

These results suggested that ParA_pSM_ ATP turnover is fine-tuned by the structural arrangement of the ParB_pSM_ and *parS_pSM_* within the PC. ParB_pSM_ assembles as a left-handed spiral to wrap *parS_pSM_* DNA without significantly distorting the DNA backbone geometry (Weihofen et al., 2006), implying that after ∼4 repeats, ParB_pSM_ dimers would make a full turn around *parS_pSM_*, positioning themselves on the same face of the nucleoprotein filament. Thus, the position of the 4_th_ repeats relative to that of the 1_st_ repeat might be functionally important. To test this possibility, we examined the ParA_pSM_ ATPase stimulation efficiency of *parS_pSM_* DNA fragments containing two copies of 3R sequences separated by 7-bp or 14-bp of non-consensus sequence (3R-1nc-3R and 3R-2nc-3R) (Fig. 1E). The 3R-1nc-3R-*parS_pSM_* fragments showed only slightly higher stimulation (*k_cat_* = 8.3 ± 3.5 h_-1_, N=8*) compared to the single 3R-*parS_pSM_* fragment, and the 3R-2nc-3R was functionally indistinguishable from the single 3R-*parS_pSM_* fragment (Fig. 1C). We conclude that disrupted 6R-*parS_pSM_* fragments are unable to recover the full stimulation of ParA_pSM_ ATPase activity. These findings indicate that the ParB_pSM_-ParA_pSM_ interactions differ when ParB_pSM_ dimers are bound to ≥4 contiguous heptad repeats compared to ParB_pSM_ dimers unbound to the repeats or bound to fewer or a disrupted array of heptad repeats.

### Substoichiometric concentration of *parS_pSM_* relative to ParB_pSM_ is sufficient to fully activate ParA_pSM_ ATPase

We originally assumed that for efficient ParA_pSM_ ATPase activation by the ParB_pSM_-*parS_pSM_* complex all ParB_pSM_ dimers need to be bound to *parS_pSM_* and accordingly maintained stoichiometrically excess *parS_pSM_*-sequence concentration relative to ParB_pSM_-dimers in the reaction. To test this assumption, we next changed the concentration of the 7R-*parS_pSM_* DNA fragment while keeping the concentrations of ParA_pSM_ and ParB_pSM_ both at 2 μM. Contrary to our expectation, significantly lower heptad consensus sequence concentrations of the 7R-*parS_pSM_* DNA fragment compared to ParB_pSM_ dimers (B_2_) were sufficient for ATPase activation (Fig. 1D). If the original assumption was correct, 2 μM ParB_pSM_ should have required 1 μM consensus sequence repeats (143 nM 7R-*parS_pSM_*) for full activation. According to the results of Fig. 1A, at high 7R-*parS_pSM_* concentration, half-activation by ParB_pSM_ required ∼650 nM ParB_pSM_, which would have required ∼325 nM *parS_pSM_* consensus sequence if full binding was needed. Observed half-saturation *parS_pSM_* repeat concentration was ∼130 nM (∼19 nM 7R-*parS_pSM_*) or less. Thus, assuming most of the ParB_pSM_ molecules in our preparations are active, it appears that at any given time, less than half of the ParB_pSM_ dimers need to be in complex with 7R-*parS_pSM_* to exert full ATPase activation.

### Stimulation of ParA_pSM_ release from nsDNA-carpet by ParB_pSM_ requires *parS_pSM_* DNA with four or more heptad repeats

Next, we examined how the dissociation of ParA_pSM_-ATP dimers from nsDNA is influenced by ParB_pSM_ in the presence of *parS_pSM_* DNA. For these experiments, we used a nsDNA-carpeted two-inlet flow cell observed under TIRF microscopy as described in Materials and Methods (Vecchiarelli et al. 2013) (Fig. 2A). We used a ParA_pSM_-GFP fusion protein and a ParB_pSM_-*cys* conjugated to Alexa647 to facilitate visualization of protein association-dissociation to/from the nsDNA-carpet. Stimulation of ParA_pSM_-GFP ATPase activity by ParB_pSM_ and 7R-*parS* was comparable to wild type ParA_pSM_ (Fig. 2-figure supplement 1). First, we tested the association/dissociation of individual proteins (bound at 1 μM) to/from the nsDNA-carpet. The affinity of ParB_pSM_-Alexa647 (1% labeled) to nsDNA was weak. ParB_pSM_-Alexa647 in the absence of ParA_pSM_ accumulated on the DNA-carpet reaching a density of ∼3000 ± 500 dimers/μm_2_ after 15 min of constant flow (5 μl/min) (Fig. 2-figure supplement 2A bottom). When the sample solution was switched to a buffer without protein, most ParB_pSM_ rapidly dissociated with kinetics that can be fitted to a double-exponential function (*k_off-fast_* = 5.1 ± 0.7 min_-1_ (73%), *k_off-slow_* = 0.11 ± 0.04 min_-1_, N=3; all *k_off_*s reported in this study represent apparent pseudo-first order rate constants) (Fig. 2-figure supplement 2B bottom). ParA_pSM_-GFP preincubated with ATP and Mg_2+_ reached a saturation density on the nsDNA-carpet of 4.25 (± 0.06) x10_4_ dimers/μm_2_ (N=4*), whereas in the absence of ATP, less than 1000 dimers/μm_2_ of ParA_pSM_ bound to the nsDNA-carpet (Fig. 2-figure supplement 2A top). Next, 1 μM ParA_pSM_-GFP preincubated with ATP was flowed onto the nsDNA-carpet until 5-10% of saturation density (∼4000 dimers/μm_2_), at which point the flow was switched to buffer containing ATP without ParA_pSM_-GFP. A small fraction of ParA_pSM_-GFP (less than ∼5%) dissociated within ∼1 min and the remainder dissociated slowly with *k_off_* of <0.013 min_- 1_ (N=3, Fig. 2B).

**Figure 2.**
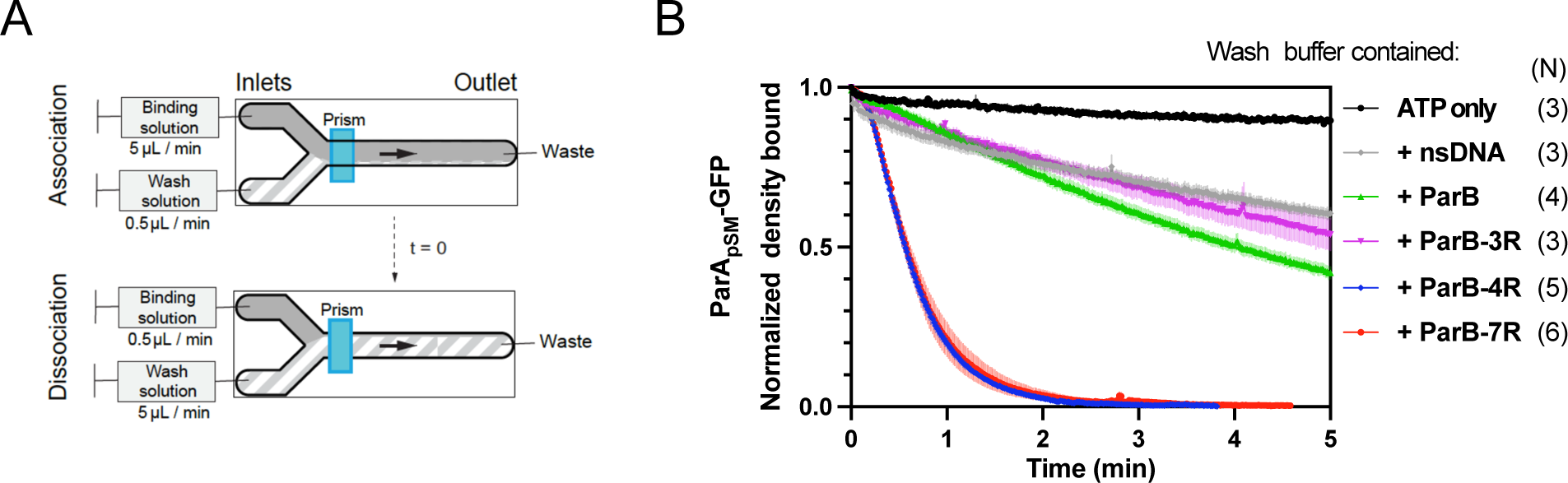
Kinetics of ParA_pSM_ disassembly from the nsDNA-carpet. (A) Schematic of two-inlet flow cell used for visualizing the association and dissociation of fluorescent proteins on nsDNA-carpet. Binding and washing solutions each containing proteins, double-stranded DNA fragments or buffer alone as specified were infused at different flow rates (5 μl/min or 0.5 μl/min) from two inlets on the left into a Y-shaped flow cell. At the middle of the flow channel just downstream of the flow convergence point where the observations are made, content of the faster infusion syringe flows over the observation point. By switching the flow rates of the two syringes, protein binding to the nsDNA-carpet and dissociation during the washing cycle can be recorded. (B) ParB_pSM_ in the presence of parS_pSM_ with at least four contiguous ParB_pSM_-binding sequence repeats stimulates the ParA_pSM_-GFP dissociation from the nsDNA-carpet. ParA_pSM_-GFP (1 μM) preincubated with 1 mM ATP was infused into the nsDNA-carpeted flow cell at 5 μl/min, while the washing solution containing the specified components was infused at 0.5 μl/min. When the ParA_pSM_-GFP density on the nsDNA-carpet reached 5-10% of the saturation density (t = 0) (∼4000 ParA**_pSM_** dimers/μm_2_), the flow rates were switched to start the wash with solution containing: buffer alone, 55-bp scramDNA fragment (65 nM in bp), or with ParB_pSM_ (1 μM) without or with different parS_pSM_ fragment (0.5 μM parS_pSM_ heptad repeat sequence). The Y-axis shows the ParA_pSM_-GFP intensity normalized to that at t = 0. Each timepoint represents the mean with error bar corresponding to the SEM of N repeated experiments. Source Data 1 Figure Supplement 1: ATPase activity of ParA_pSM_-GFP and the hydrolysis-deficient ParA_pSMD60E_-GFP. Figure Supplement 2: ParA_pSM_, ParA_pSMD60E_ and ParB_pSM_ interactions with the DNA-carpet measured each separately. Figure Supplement 3: ParB_pSM_-activated dissociation of ParA_pSM_-GFP or ParA_pSMD60E_-GFP bound to nsDNA- carpet to saturation density.

The addition of competitor nsDNA (71 nM scrambled 55-bp duplex DNA) in the wash buffer slightly increased the *k_off_* of majority (∼93%) fraction of ParA_pSM_-GFP (*k_off_* = 0.075 ± 0.001 min_-1_, N=3), perhaps in part by competing with rebinding of ParA_pSM_-GFP-ATP to the nsDNA-carpet (Fig. 2B). When 1 μM ParB_pSM_ (unlabeled or 1% Alexa647 labeled) was added to the wash buffer without competitor nsDNA, *k_off_* of ParA_pSM_-GFP from nsDNA-carpet reached 0.17 ± 0.01 min_-1_ (N=4, Fig. 2B). ParA_pSM_-GFP dissociation from the nsDNA-carpet was accelerated when wash solution contained 71 nM 7R-*parS_pSM_* DNA (0.5 μM of the consensus sequence repeat) and 1 μM ParB_pSM_ (*k_off_* = 1.97 ± 0.09) after a brief period of initial slower dissociation rate (see below for details) (Fig. 2B). This was ∼12- or ∼26-fold higher *k_off_* compared to ParB_pSM_ or nsDNA wash, respectively. Similarly, 4R-*parS_pSM_* DNA supported accelerated ParA_pSM_-GFP release by ParB_pSM_, but 3R-*parS_pSM_* DNA did not (Fig. 2B). The results paralleled those shown in Figure 1A solidifying the notion that a functional *parS_pSM_* must carry four copies of ParB_pSM_ dimer binding sequence repeats.

The dependence of ParA_pSM_ dissociation behaviors on the heptad repeat number of the *parS_pSM_* fragment combined with 1 μM ParB_pSM_ in the wash solution when the wash was started with ParA_pSM_ bound to the nsDNA-carpet at a saturating density were qualitatively similar. However, the dissociation kinetics were significantly slower (Fig. 2-figure supplement 3A), likely reflecting the ∼10-fold higher starting ParA_pSM_-GFP density on the nsDNA-carpet, for which the concentration of ParB_pSM_ used likely was limiting.

### Accelerated release of ParA_pSM_ from DNA-carpet starts only after accumulation of ParB_pSM_ on the carpet in the presence of 7R- or 4R-*parS_pSM_*

The accelerated ParA_pSM_-GFP release from the nsDNA-carpet when washed with ParB_pSM_-4R or -7R *parS_pSM_* complexes started after an initial slower rate of ParA_pSM_ release (Fig. 2B). This time lag prompted us to examine the binding kinetics of ParB_pSM_ to the ParA_pSM_-bound nsDNA-carpet at different concentrations of ParB_pSM_. Upon switching the flow to the washing solution without *parS_pSM_*, ParB_pSM_ bound to the ParA_pSM_-bound nsDNA-carpet to a peak density of **∼**3000 dimers/μm_2_ and then quickly decreased to a plateau density of **∼**2000 dimers/μm_2_ (Fig. 3A-d). The initial overshoot of ParB_pSM_ binding appeared to roughly coincide with the fast dissociation phase of ParA_pSM_. During the subsequent slow dissociation of ParA_pSM_, ParB_pSM_ density on the nsDNA- carpet remained relatively constant with the protein ratio reaching ∼1 after several min of washing with 1 μM ParB_pSM_ (Fig. 3A-d).

**Figure 3.**
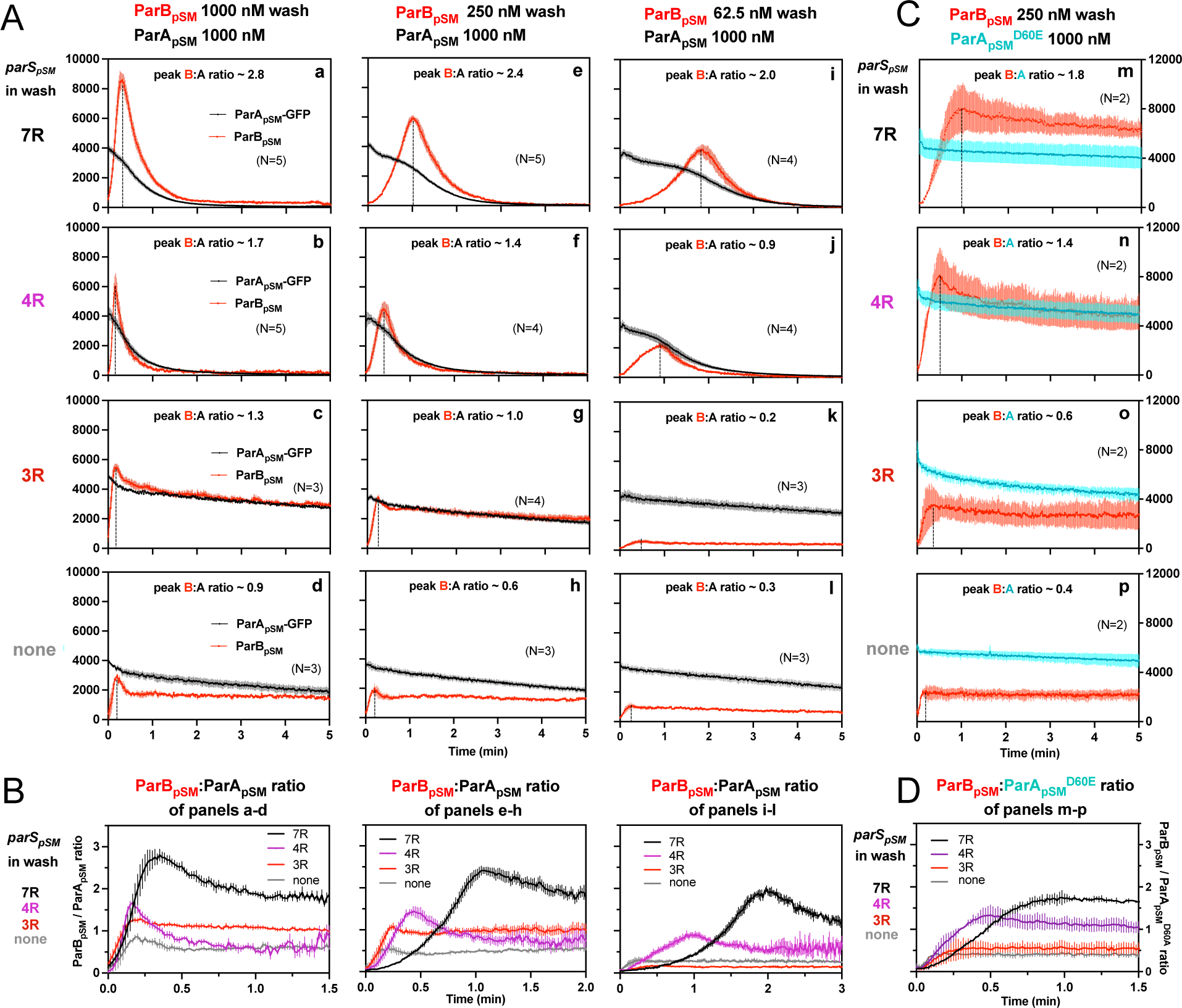
ParB_pSM_-*parS_pSM_* concentration affects kinetics of ATP hydrolysis-dependent ParA_pSM_ disassembly from the nsDNA-carpet. (A) ParA_pSM_-GFP (1 μM) preincubated with 1 mM ATP was infused into the nsDNA-carpeted flow cell at 5 μl/min and when the ParA_pSM_-GFP density on the nsDNA- carpet reached ∼10% of the saturation density, the flow over the observation area was switched (t = 0) to wash solution containing ParB_pSM_ (62.5, 250, or 1000 nM) and the stoichiometric concentration of parS_pSM_ fragments indicated in the “parS_pSM_ in wash” column on the left side of each row of panels. Fluorescence signal was converted to protein density and plotted (dimers/μm_2_, ParA_pSM_-GFP: black and ParB_pSM_-Alexa647: red). Each time point represents mean and SEM (vertical spread) of N repeated experiments. Dashed vertical lines indicate the peak ParB_pSM_ density on the nsDNA-carpet. (B) The time course of the ParB_pSM_:ParA_pSM_ molar ratio (B/A) for the four panels in the columns in A with matching ParB_pSM_ concentration with different parS_pSM_ (7R: black, 4R: purple, 3R: red, none: gray) in the wash solution. (C) ATP hydrolysis is required for accelerated ParA_pSM_ release from the nsDNA-carpet. The experiments shown in A (middle column) were repeated using ParA_pSMD60E_- GFP (1 μM) bound to the nsDNA-carpet and ParB_pSM_ (250 nM) plus parS_pSM_ fragment (125nM heptad concentration) in the wash solution. Fluorescence signal was converted to protein density (dimers/μm_2_, ParA_pSMD60E_-GFP: cyan and ParB_pSM_-Alexa647: red) and plotted. Each time point represents mean and SEM (vertical spread) of N experiments. (D) The time course of the ParB_pSM_:ParA_pSMD60E_ molar ratio (B/A_D60E_) for the four panels of C (7R: black, 4R: purple, 3R: red, none: gray). Each time point represents the mean with error bar corresponding to the SEM of N repeated experiments. Source Data 1

In the presence of 1 μM ParB_pSM_ and 7R-*parS_pSM_* in the wash solution, ParA_pSM_-GFP release from the nsDNA-carpet took place in two phases (Fig. 3A-a). First, ParB_pSM_ accumulated on the ParA_pSM_-bound nsDNA-carpet to a density two-fold or more in excess of the carpet-bound ParA_pSM_. During this phase, ParA_pSM_ dissociated from the nsDNA-carpet relatively slowly. Next, as the binding of ParB_pSM_ stopped and began to quickly dissociate (*k_off_* = 2.4 ± 0.1 min_-1_, N=5), ParA_pSM_ dissociation also accelerated to a higher rate (*k_off_* = 1.8 ± 0.05 min_-1_, N=5) (Fig. 3A-a). Near-complete dissociation of both proteins from the nsDNA-carpet occurred within a few minutes. Biphasic dissociation kinetics of ParA_pSM_ was clearer when the wash solution contained lower concentrations of the 7R-*parS_pSM_*-ParB_pSM_ complex, as ParB_pSM_ accumulated on the ParA_pSM_- bound nsDNA-carpet slower, and it took longer to transition to the accelerated dissociation phase (Fig. 3A-e, 3A-i). This indicates that the initial nsDNA-carpet-bound ParA_pSM_-ParB_pSM_ complex is not activated for ParA_pSM_ dissociation from nsDNA, and slow transition of the nsDNA-bound complex is necessary to start the ParA_pSM_ disassembly.

We next examined the ability of 4R- and 3R-*parS_pSM_* DNA to accelerate ParA_pSM_-GFP dissociation in the presence of ParB_pSM_. Switching 7R-*parS_pSM_* in the wash solution to 4R-*parS_pSM_* (1 μM ParB_pSM_, 125 nM 4R DNA) yielded qualitatively similar two-step dissociation curves (Fig. 3A-b). The *k_off_* of ParA_pSM_ for the accelerated dissociation phase (1.9 ± 0.06 min_-1_, N=5) was roughly the same as in the presence of 7R-*parS_pSM_*. After the peak of binding, ParB_pSM_ dissociated from the nsDNA-carpet with *k_off_* of 3.9 ± 0.2 min_-1_, N=5. The dissociation burst of ParA_pSM_-GFP was triggered in the presence of 4R-*parS_pSM_* at ParB_pSM_/ParA_pSM_ molar ratio of ∼1.7, compared to ∼2.8 in the presence of 7R-*parS_pSM_* (at 1 μM ParB_pSM_). This difference likely in part reflects the fact that an individual ParB_pSM_-*parS_pSM_* complex with 4R-*parS_pSM_* contains fewer ParB_pSM_ dimers than the 7R-*parS_pSM_* complex contains. Also, at a given ParB_pSM_ concentration in the wash solution, washing with 7R-*parS_pSM_* took roughly twice as long to reach the peak ParB_pSM_/ParA_pSM_ ratio to start the rapid ParA_pSM_ dissociation phase compared to the 4R-*parS_pSM_* wash (Fig. 3B). We believe this likely reflects, in part, a lower concentration of the larger ParB_pSM_-7R-*parS_pSM_* complex than the complex with 4R-*parS_pSM_*. The accelerated phase of ParA_pSM_ *k_off_* appeared higher in the presence of higher concentrations of 7R- or 4R-*parS_pSM_*-ParB_pSM_ complex in the wash solution (Table 1).

**Table 1.** The accelerated ParA_pSM_ dissociation phase of the curves in Fig. 3A-a, -e, -i and 3A-b, -f, -j after the peak ParB_pSM_/ParA_pSM_ ratio points were fitted to single exponential curves to estimate the k_off_ values. The means with error bars corresponding to the SEM of N repeated experiments in Figure 3 are shown.

|  | 7R-*parS<sub>pSM</sub>* | 4R-*parS<sub>pSM</sub>* |
| --- | --- | --- |
| [ParB<sub>pSM</sub>] nM | ParA<sub>pSM</sub> $k_{off}$ | ParA<sub>pSM</sub> $k_{off}$ |
| 1000 | 1.808 ± 0.045 | 1.909 ± 0.057 |
| 250 | 1.495 ± 0.024 | 1.462 ± 0.040 |
| 62.5 | 1.184 ± 0.041 | 1.101 ± 0.033 |

Unlike the results above using 7R- or 4R-*parS_pSM_* in the wash solution, washing experiments with 3R-*parS_pSM_* (1 μM ParB_pSM_, 166 nM 3R-*parS_pSM_*) did not exhibit an accelerated ParA_pSM_-GFP dissociation phase, as in the absence of *parS_pSM_* (Fig. 3A-c, -d). ParB_pSM_ quickly accumulated onto the ParA_pSM_-bound nsDNA-carpet up to a ParB_pSM_/ParA_pSM_ ratio of ∼1.3 (1 μM ParB_pSM_). Thus, ParB_pSM_ appears to be able to associate with nsDNA-bound ParA_pSM_ to a stoichiometry which is unlikely to be the factor preventing stimulation of ParA_pSM_ dissociation. Both ParA_pSM_-GFP and ParB_pSM_ dissociated from their peak density on the carpet with double-exponential kinetics roughly in parallel (ParA_pSM_-GFP; *k_off-fast_* = 3.8 min_-1_ ± 0.4 min_-1_ (∼19%), *k_off-slow_* = 0.07 ± 0.003 min_-1_: ParB_pSM_; *k_off-fast_* = 1.4 ± 0.3 min_-1_ (∼24%), *k_off-slow_* = 0.05 ± 0.004 min_-1_, N=3) (Fig. 3A-c).

### ATP hydrolysis triggers fast ParA_pSM_ dissociation from the nsDNA-carpet

We next asked if the accelerated dissociation of ParA_pSM_ from the nsDNA-carpet in the presence of *parS_pSM_* fragments depends on ATP hydrolysis by ParA_pSM_. Among ParA_pSM_ homologues, a conserved Asp in the Walker A’ motif is critical for ATP hydrolysis (Pratto et al. 2008, Park et al. 2012). We prepared the ParA_pSMD60E_-GFP fusion protein; it exhibited no significant residual ATPase activity and faster ATP-dependent nsDNA binding compared to the wild-type protein (Fig. 2-figure supplement 1, Fig. 2-figure supplement 2A top). The apparent *k_off_* of the majority fraction (>80%) of nsDNA-bound ParA_pSMD60E_-GFP washed with ATP-containing buffer (0.015 ± 0.007 min_-1_, N=2) was not significantly different from ParA_pSM_-GFP (0.014 ± 0.002 min_-1_, N=2) (Fig. S5B top). However, when the wash solution contained 250 nM ParB_pSM_ and 36 nM 7R-*parS_pSM_*, ParA_pSMD60E_-GFP dissociation from the nsDNA-carpet was only a factor of ∼2 faster (Fig. 3C-a; *k_off_* **=** 0.029 ± 0.005 min_-1_, N=2) than washing with ATP-buffer. As washing started, the ParB_pSM_ associated with the carpet-bound ParA_pSMD60E_-GFP to a density well beyond that of ParA_pSMD60E_- GFP with similar association kinetics as with the carpet-bound ParA_pSM_-GFP (Fig. 3A-e). The ParB_pSM_/ParA_pSM_ ratio bound to the nsDNA-carpet at the initial ParB_pSM_ overshoot peak was ∼1.75 in the presence of 7R-*parS_pSM_*, ∼1.3 with 4R-*parS_pSM_* and ∼0.6 with 3R-*parS_pSM_* (Fig. 3D). After the peak density, ParB_pSM_ dissociated from the nsDNA-carpet slowly (*k_off_* = 0.036 ± 0.003 min_-1_, N=2) (Fig. 3C-a). Thus, the accelerated DNA dissociation of ParA_pSM_ by the active ParB_pSM_- *parS_pSM_* complex is coupled to ATP hydrolysis, and in the absence of hydrolysis, ParA_pSMD60E_- ATP dimers stayed stably on nsDNA while associated with active *parS_pSM_*-ParB_pSM_ complex.

### ParB_pSM_ associates with nsDNA-bound ParA_pSM_ more stably in the presence of 7R- or 4R-***parS_pSM_* prior to ATP hydrolysis**

In the experiments of Figures 2 and 3, ParA_pSM_-ATP dimers were bound to the nsDNA-carpet first without ParB_pSM_, and during the wash phase of the experiments, the carpet-bound ParA_pSM_ was constantly exposed to a solution containing ParB_pSM_ and *parS_pSM_*. Because ParB_pSM_ and *parS_pSM_* dissociating from carpet-bound ParA_pSM_ could be exchanged by those in the wash solution, the data did not report the stability of ParA_pSM_-ParB_pSM_ interaction on the nsDNA-carpet. In the next set of experiments, we pre-incubated an equimolar ratio of ParA_pSM_-GFP (1μM), ParB_pSM_ (1 μM), and 7R-, 4R-, or 3R-*parS_pSM_* (0.5 μM total concentration of the consensus sequence repeats) in the presence of ATP for 30 min at room temperature. The samples were then infused into the nsDNA-carpeted flow cell at 5 μl/min for 15 min. At this point, the incoming components interacting with the nsDNA-carpet would be approaching a steady state. The carpet-bound protein complexes were then washed with a buffer containing ATP without ParA_pSM_, ParB_pSM_, or *parS_pSM_*. ParA_pSM_-GFP- ATP in the absence of ParB_pSM_ reached saturation on the nsDNA-carpet in 5 to 10 min and then was released from the carpet slowly during the wash phase with dissociation kinetics that fit a double exponential function, majority fraction dissociating slowly (*k_off-fast_* = 1.7 ± 1.1 min_-1_ (∼7%), *k_off-slow_* = 0.014 ± 0.002 min_-1_, N=2) (Fig. 2-figure supplement 2B top) as observed starting with lower ParA_pSM_-GFP density (Fig. 2B).

When ParA_pSM_-GFP was mixed with ParB_pSM_ and ATP in the absence of *parS_pSM_* and infused into the flow cell, ParA_pSM_-GFP-ATP reached near saturation density on the nsDNA-carpet in ∼10 min. During the wash phase, ParA_pSM_-GFP dissociated from the nsDNA-carpet with double exponential kinetics (*k_off-fast_* = 2.3 ± 0.5 min_-1_, *k_off-slow_* = 0.057 ± 0.01 min_-1_, N=4) as in the absence of ParB_pSM_ except for the several fold faster slow phase *k_off_* and increased fraction of faster-dissociating population (∼30%) (Fig. 4A-d). When the reaction mixture also contained 3R-*parS_pSM_*, ParA_pSM_- GFP dissociation kinetics were similar to the above, except reduced fraction of the faster dissociating ParA_pSM_-GFP population (Fig. 4A-c). In these experiments, ParB_pSM_ first accumulated on the nsDNA-carpet to roughly two-thirds of the density of ParA_pSM_. When washing started, the majority (∼75%) of ParB_pSM_ dissociated quickly (∼4 or ∼7 min_-1_, with or without 3R-*parS_pSM_*). Thus, even in the presence of 3R-*parS_pSM_* fragment, most ParB_pSM_ dissociates from ParA_pSM_ bound to nsDNA-carpet within half of a minute (Fig. 4A-c, 4B).

**Figure 4.**
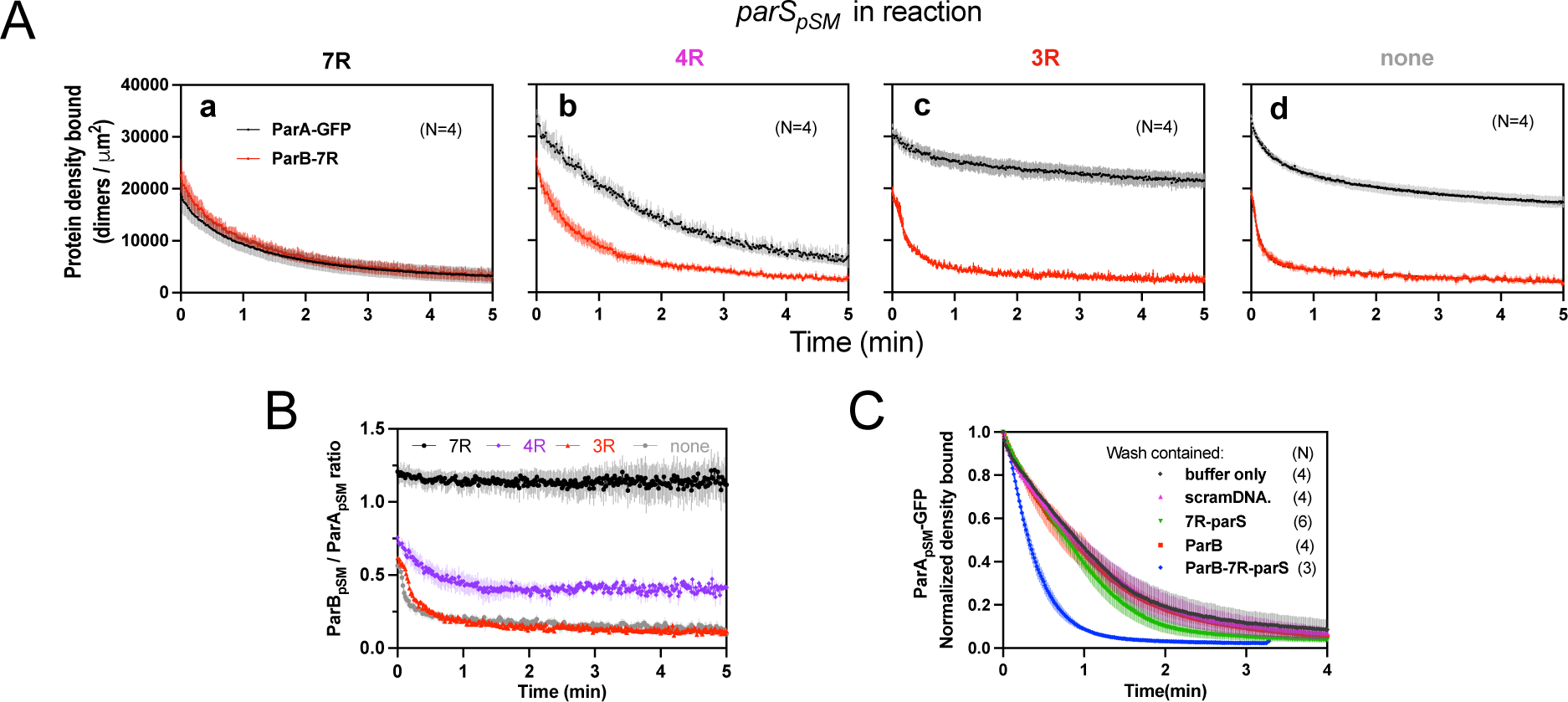
Stable ParA_pSM_-ParB_pSM_ complex is formed prior to ATP hydrolysis in the presence of functionally active *parS_pSM_*. (A) ParA_pSM_-GFP (1μM) and ParB_pSM_-Alexa647 (1% labelled, 1μM) were preincubated with ATP (1mM) plus 7R-, 4R-, 3R-*parS_pSM_*, or without *parS_pSM_.* The preincubated sample was infused into the nsDNA-carpeted flow cell at 5μl/min for 15 min and then the solution flowing over the observation area was switched to a buffer containing ATP (t = 0). Fluorescence signals of ParA_pSM_ (black) and ParB_pSM_ (red) were converted to protein density on the nsDNA-carpet (dimers per μm_2_) and plotted. Each time point represents mean and SEM (vertical spread) of N experiments. (B) Time course of the carpet-bound ParB_pSM_:ParA_pSM_ molar ratio for the four panels of A (7R: black, 4R: purple, 3R: red, none: gray). (C) Continued supply of ParB**_pSM_** and *parS_pSM_* in the washing solution is required to sustain high rate of ParA_pSM_ disassembly. The experiment in the presence of 7R-*parS_pSM_* in panel A-a was repeated with addition of ParB**_pSM_** and/or 7R- *parS_pSM_* in the wash solution. Fluorescence signals of ParA_pSM_ was normalized to the value at t = 0. Each time point represents mean and SEM (vertical spread) of N experiments. Source Data 1

When ParA_pSM_-GFP and ParB_pSM_ were preincubated with 7R-*parS_pSM_*, the steady state ParA_pSM_ accumulation on the nsDNA-carpet was suppressed by nearly 50%, and ParB_pSM_ and ParA_pSM_ accumulated on the nsDNA-carpet at approximately ∼1A:1.2B ratio. When the wash was started, the two proteins dissociated from the carpet maintaining roughly constant protein stoichiometry with kinetics that can be fit to a double-exponential curve (*k_off-fast-A_* (∼73%) 0.92 ± 0.18 min_-1_, *k_off- slow-A_* 0.075 ± 0.04 min_-1_, and *k_off-fast-B_* (∼70%) 1.1 ± 0.14 min_-1_, *k_off-slow-B_* 0.13 ± 0.03 min_-1_, N=4; Fig. 4A-a). Thus, before ATP hydrolysis disassembles ParA_pSM_ dimer from the nsDNA-carpet, ParA_pSM_-ParB_pSM_ protein interaction appears to be significantly more stable in the presence of 7R- *parS_pSM_* compared to the complex involving 3R-*parS_pSM_* (Fig. 4B).

In the presence of 4R-*parS_pSM_*, carpet binding and dissociation dynamics of the two proteins were qualitatively similar to those in the presence of 7R-*parS_pSM_*, except for the following differences: 4R-*parS_pSM_* did not suppress ParA_pSM_ binding to the nsDNA-carpet as effectively as 7R-*parS_pSM_*, the ParB_pSM_/ParA_pSM_ ratio at the end of 15 min sample infusion was ∼0.75, which dropped to ∼0.5 within one min of washing after which, the ratio remained constant. Observed *k_off_* were: ParA_pSM_, (*k_off-fast-A_* (60%) 0.67 ± 0.14 min_-1_, *k_off-slow-A_* 0.17 ± 0.05 min_-1_; and *k_off-fast-B_* (66%) 1.9 ± 0.2 min_-1_, *k_off-slow-B_* 0.23 ± 0.02 min_-1_, N=4; Fig. 4A-b). It appears that ParA_pSM_ bound to the nsDNA-carpet interacts slightly less stably with ParB_pSM_ in the complex assembled with 4R-*parS_pSM_* compared to those assembled with 7R-*parS_pSM_*, but it was still significantly more stable compared to the majority of the ParA_pSM_-ParB_pSM_ complexes that accumulate in the presence of 3R-*parS_pSM_* or without *parS_pSM_* DNA.

In this experiment, the absence of ParB_pSM_ and 7R- or 4R-*parS_pSM_* in the wash solution flowing over the nsDNA-carpet appeared to have caused a slowdown of dissociation of the ParA_pSM_- ParB_pSM_ complexes from nsDNA compared to the experiments of Figures 2B and 3A. This prompted us to examine if partial loss of the *parS_pSM_* DNA and/or ParB_pSM_ from the complex during the washing caused this kinetic change. We repeated the experiments of Figure 4A-a with the addition of ParB_pSM_ (1 μM) and/or 7R-*parS_pSM_* DNA fragment (500 nM consensus repeats) in the washing solution. The addition of ParB_pSM_ and 7R-*parS_pSM_* in the washing solution significantly accelerated ParA_pSM_ *k_off_* to 2.5 ± 0.25 min_-1_ (N=3), a slightly higher level than seen in Figure 3A-a (1.8 ± 0.05 min_-1_), which might have been an underestimate due to the preceding slow complex assembly steps. The addition of ParB_pSM_ or 7R-*parS_pSM_* separately to the washing solution did not have significant effects on the ParA_pSM_ dissociation kinetics (Fig. 4C). Thus, partial loss of ParB_pSM_-*parS_pSM_* complex from the nsDNA-bound ParA_pSM_ at the onset of the wash (7R- and 4R-*parS_pSM_*) and also during the wash (4R-*parS_pSM_*) appears to have compromised the efficiency of ParA_pSM_ dissociation from nsDNA in the experiments of Figure 4A-a, and -b. This indicates that not all the ParB_pSM_-*parS_pSM_* complexes that can contribute to dissociation of ParA_pSM_ from nsDNA are stably retained *via* ParA_pSM_ to the nsDNA-carpet through the course of reaction.

### ParA_pSM_-ATP in the wash solution stabilizes functional ParB_pSM_*-parS_pSM_* complexes associated with carpet-bound ParA_pSM_

When preformed ParA_pSM_-ParB_pSM_ complexes bound to the nsDNA-carpet in the presence of 7R- or 4R-*parS_pSM_* DNA were washed with buffer containing ATP-activated ParA_pSM_-GFP (0.5μM), freshly arriving ParA_pSM_-ATP dimers on the nsDNA-carpet replacing the dissociating ParA_pSM_ partially stabilized ParB_pSM_ (Fig. 5-a, -b). This observation indicates that when ParA_pSM_ dissociates from nsDNA following ATP hydrolysis, some fraction of ParB_pSM_ in the presence of 7R- or 4R- *parS_pSM_* that induced ATP hydrolysis, can capture the newly arriving ParA_pSM_-ATP dimers on the nsDNA-carpet. The complex containing 7R-*parS_pSM_* was more efficient for recapture than the 4R- *parS_pSM_* complex. We propose larger numbers of ParB_pSM_ dimers complexed with *parS_pSM_* with more sequence repeats can interact simultaneously with multiple ParA_pSM_-nsDNA mini-filaments, more readily capturing newly arrived ParA_pSM_-ATP dimers at nearby nsDNA sites. Evidence supporting such multivalent interactions between nsDNA-bound ParA_pSM_ and *parS_pSM_*-bound ParB_pSM_ has been reported previously (Pratto et al., 2009). ParB_pSM_ stabilization by replenishment of ParA_pSM_-ATP was not observed in the presence of 3R-*parS_pSM_* or in the absence of *parS_pSM_*, consistent with the notion that ParA_pSM_-ParB_pSM_ interaction under these conditions are unstable (Fig. 5-c, -d).

**Figure 5.**
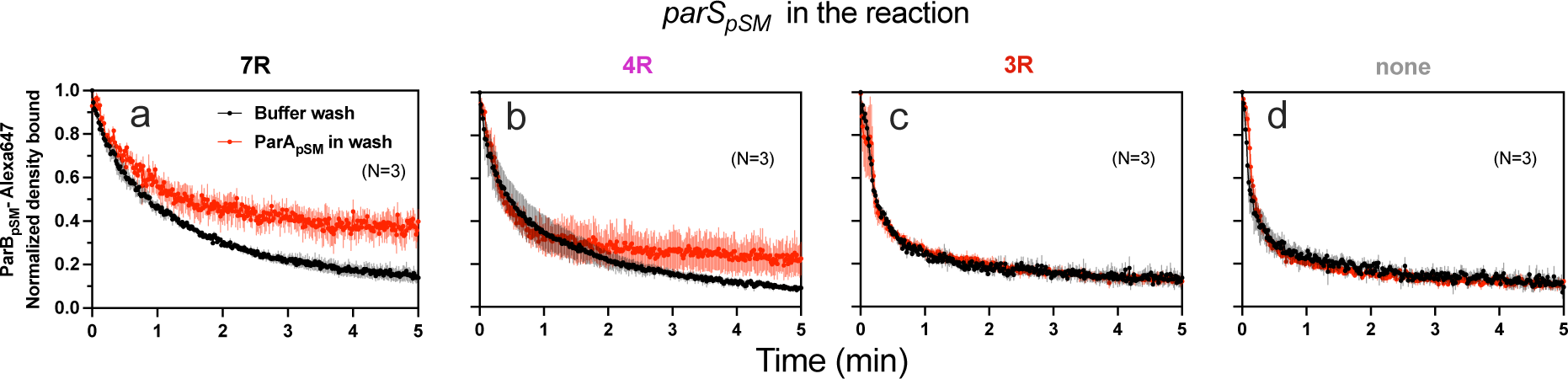
ParA_pSM_-ATP stabilizes functionally active *parS_pSM_*-bound ParB_pSM_ on the nsDNA- carpet. Complexes containing ParA_pSM_-GFP, and ParB_pSM_-Alexa647 bound to the DNA-carpet together with different *parS_pSM_* DNA fragments were washed with either a buffer containing 1mM ATP (black) or the same buffer containing ParA_pSM_-GFP (0.5 μM, red), in addition to ATP. The ParB_pSM_-Alexa647 dissociation curves normalized to the protein density at t = 0 were plotted. Each time point represents mean and SEM (vertical spread) of N experiments. Source Data 1

## Discussion

Using a combination of steady-state ATPase assays and nsDNA dissociation kinetic measurements of ParA_pSM_-ParB_pSM_ complexes in the presence of a variety of modified *parS_pSM_*-DNA fragments, we studied the molecular requirements for activation of ParA_pSM_. This work revealed distinct *parS_pSM_* structural requirements for efficient activation of ParA_pSM_ ATPase by ParB_pSM_ and a multi-step assembly process for the active *parS_pSM_*-ParB_pSM_-ParA_pSM_ complex. The *parS_pSM_* DNA must contain at minimum four contiguous consensus sequence repeats to enable ParB_pSM_ dimer binding in a state that induces ATP hydrolysis and accelerated dissociation of ParA_pSM_ from nsDNA. The structural requirements for *parS_pSM_* function assure high specificity of the centromere site for PC function.

### ParB_pSM_-*parS_pSM_* recognition by ParA_pSM_

The *parS_pSM_* sequence features required for the function of ParB_pSM_, an RHH-ParB protein contrast with those for the HTH-ParB proteins. Whereas natural *parS_F_* sequence for the F-plasmid is composed of 12 repeats of the ParB_F_ dimer binding sequence, a single copy of this consensus sequence is able to support faithful plasmid partition (Biek and Shi 1994). Further, HTH-ParB proteins can spread around *parS* sites, forming large PCs containing many ParB molecules, most of which have moved away from *parS* sequence. In contrast, RHH-ParB proteins lack the CTPase domain that is critical for HTH-ParB spreading (Soh et al. 2019, Jalai et al. 2020, Osorio-Valeriano et al. 2019) and they have not been shown to spread from *parS* sites. Systems involving RHH- ParB proteins perhaps evolved a fundamentally different system architecture of PC dynamics to accomplish robust partitioning of replicated plasmid copies. It is currently unclear what constitutes the defining feature of the ParB_pSM_ molecules bound to four or more contiguous *parS_pSM_* heptad sequence repeats: we offer a possible scenario below.

### Assembly/disassembly dynamics of ParAB_pSM_ complex on the nsDNA

ParA_pSM_-ATP is slow to dissociate from the nsDNA-carpet in the presence of ParB_pSM_ alone, or ParB_pSM_-3R-*parS_pSM_* complex in the wash solution. We showed that ParB_pSM_ can quickly interact with nsDNA-bound ParA_pSM_ in the absence of functional *parS_pSM_* DNA (Fig. 3A-c). However, this ParA_pSM_-ParB_pSM_ interaction does not fully activate the ATPase (Fig. 1). All wash curves of ParA_pSM_-ATP dimers bound to nsDNA-carpet under conditions that do not trigger efficient ATPase activation exhibited double-exponential dissociation kinetics, although the small amplitude of the fast dissociation phase, typically ∼5%, made quantitative comparison difficult (Fig. 2B, 3A). The fast dissociation phase had a time scale of less than 1 min, while the majority fraction dissociated much slower under conditions that do not activate the ATPase. This indicated that the carpet-bound ParA_pSM_ dimers were composed of two distinct state populations, transitions between which are slow. ParB_pSM_ with or without 3R-*parS_pSM_*, after the initial rapid binding to nsDNA-bound ParA_pSM_, also dissociated with double-exponential kinetics; a fraction of the carpet- associated ParB_pSM_ quickly dissociated within 1 min before settling to a quasi-steady state with the constant supply of ParB_pSM_ or ParB_pSM_-3R-*parS_pSM_* complex in the wash solution and slowly dissociating ParA_pSM_ on the nsDNA-carpet (Fig. 3A-c, d). We speculate that this small initial binding overshoot reflects ParB_pSM_ association to the less populated fast dissociating fraction of the carpet-bound ParA_pSM_ dimers, which presumably are in a state with faster ParB_pSM_-association rate, lower nsDNA affinity, and perhaps closer to the ATP hydrolysis-competent state.

When ParA_pSM_-ATP-ParB_pSM_ complexes bound to the nsDNA-carpet in a steady state with or without 3R-*parS_pSM_* was washed with a simple buffer (Fig. 4A-c, d), a double-exponential dissociation of ParA_pSM_ was again observed with increased fraction (up to ∼30%) of the fast- dissociating population. This supports the notion that the fast-dissociating ParA_pSM_ population is in a state closer to, but not committed to, ATP hydrolysis. We hypothesize this ParA_pSM_ transitory state becomes more populated during pre-incubation with ParB_pSM_ on the nsDNA-carpet accounting for the moderate stimulation of the DNA-bound ParA_pSM_ ATPase by ParB_pSM_ or ParB_pSM_-3R-*parS_pSM_* complex at steady-state (Fig. 1A). However, this “fast” dissociation does not depend on ATP hydrolysis, considering the ATPase defective mutant ParA_pSMD60E_ also exhibits clear double-exponential dissociation (Fig. 2-figure supplement 2B top, Fig. 2-figure supplement 3B).

When ParA_pSM_-ATP prebound to nsDNA-carpet in the flow cell was washed with a solution containing fully functional 7R-*parS_pSM_*-ParB_pSM_ complexes, accelerated release of ParA_pSM_ from nsDNA was observed (Fig. 2B and Fig. 3A-a, -e, -i). However, the initial ParA_pSM_ dissociation kinetics resembled the ParB_pSM_ wash without fully active *parS_pSM_* and accelerated disassembly started with a delay. This clear transition of the dissociation mode indicates the initial complex between the nsDNA-bound ParA_pSM_ and *parS_pSM_*-bound ParB_pSM_ does not immediately trigger ATP hydrolysis. Rather a series of steps must take place subsequent to the initial association of the two protein-DNA complexes to trigger ATP hydrolysis and complex disassembly. This process appears to involve participation of additional ParB_pSM_ dimers and/or *parS_pSM_* beyond the initial complex, considering that the delay time before the accelerated ParA_pSM_ dissociation phase depends on their concentration in the wash solution while clear biphasic kinetic feature was observed even at limiting ParB_pSM_ concentrations.

The ParA_pSM_-ParB_pSM_ interaction is not stable in the absence of 7R- or 4R- *parS_pSM_* (Fig. 4B). In contrast, interaction between nsDNA-bound ParA_pSM_ and ParB_pSM_ in the presence of 7R*-parS_pSM_* is significantly more stable prior to ATP hydrolysis-dependent dissociation of ParA_pSM_ from nsDNA (Fig. 4A-a). Thus, ParA_pSM_-ATP interacts with ParB_pSM_ in the presence of fully active *parS_pSM_* in a distinct manner than in its absence. For simplicity, we propose this transition to stably interacting ParA_pSM_-ParB_pSM_ complex bound to nsDNA is the limiting step necessary before ATP hydrolysis-dependent acceleration of ParA_pSM_ dissociation. Slower *k_off_* of ParA_pSM_ from nsDNA in the experiments of Fig. 4-a and -b compared to those in Figure 2B and Fig. 3A-a and -b, and recovery of faster *k_off_* by the replenishment of ParB_pSM_ and *parS_pSM_* in the wash solution (Fig. 4C) indicated that ParA_pSM_-ParB_pSM_ complex must turn over to maintain high ParA_pSM_ *k_off_*. This suggests that not all the nsDNA-bound ParA_pSM_ dimers are interacting with ParB_pSM_ in the state committed for ATPase activation in the initial set of active complexes that form on the nsDNA- carpet.

### A model for ParB_pSM_ activation of ParA_pSM_-ATPase

Without further experimental constraints, our consideration of the mechanism of ParA_pSM_-ATPase activation by ParB_pSM_ remains speculative. Unlike other members of ParA-family of ATPases studied before, such as ParA_F_ (Taylor et al. 2021) or MinD (Vecchiarelli et al. 2016), simple interaction of ParB_pSM_-N-terminal domain, ParB_pSM1-27_, with ParA_pSM_ dimers did not efficiently activate the ATPase (Fig. 1-figure supplement 1B). Because of the helical nature of the ParB_pSM_- *parS_pSM_* complex (Weihofen et al. 2006) and ParA_pSM_-nsDNA mini-filament formation (Pratto et al. 2009), two ParA_pSM_ dimers within a mini-filament might be approximately in position to interact with ParB_pSM_ dimers separated by two intervening *parS_pSM_* sequence repeats on one face of the helical filament. Let us assume, however, that the pitch and/or the angular arrangements of these two pairs of protein dimers are not in perfect match allowing only partial interaction between ParA_pSM_ dimers and ParB_pSM_ dimers in the basal state structures of the two mini-filaments (Fig. 6A). Then, the establishment of divalent interactions and full engagement between the two protein-DNA complexes perhaps would force distortions of the structures of both of the protein-DNA mini-filaments (Fig. 6B). The force imposed upon ParA_pSM_-mini-filament might act as a steppingstone for the conformational change of ParA_pSM_ necessary for ATPase activation. Such a scenario explains the requirement for the fourth *parS_pSM_* consensus sequence repeat for the ATPase activation. We propose the specific mechanical properties of the contiguous ParB_pSM_-*parS_pSM_* mini-filament, likely different from gapped mini-filaments, is required for efficient ATPase activation, explaining the requirement for the *parS_pSM_* sequence repeat contiguity. The divalent interactions between one ParB_pSM_-*parS_pSM_* mini-filament with a ParA_pSM_-nsDNA mini-filament, with a non-interacting middle segment separating the two interacting pairs, might allow the ParB_pSM_-*parS_pSM_* complex to semi-processively activate multiple ParA_pSM_-nsDNA complexes in succession by an inchworm-like transfer to a new ParA_pSM_-nsDNA mini-filament. The ability of the otherwise dissociating 7R-*parS_pSM_*-ParB_pSM_ complex after ATPase activation and dissociation of one partner ParA_pSM_ mini-filament to recapture a freshly arriving ParA_pSM_-ATP dimers on the nsDNA-carpet (Fig. 5-a) is consistent with such a possibility.

**Figure 6.**
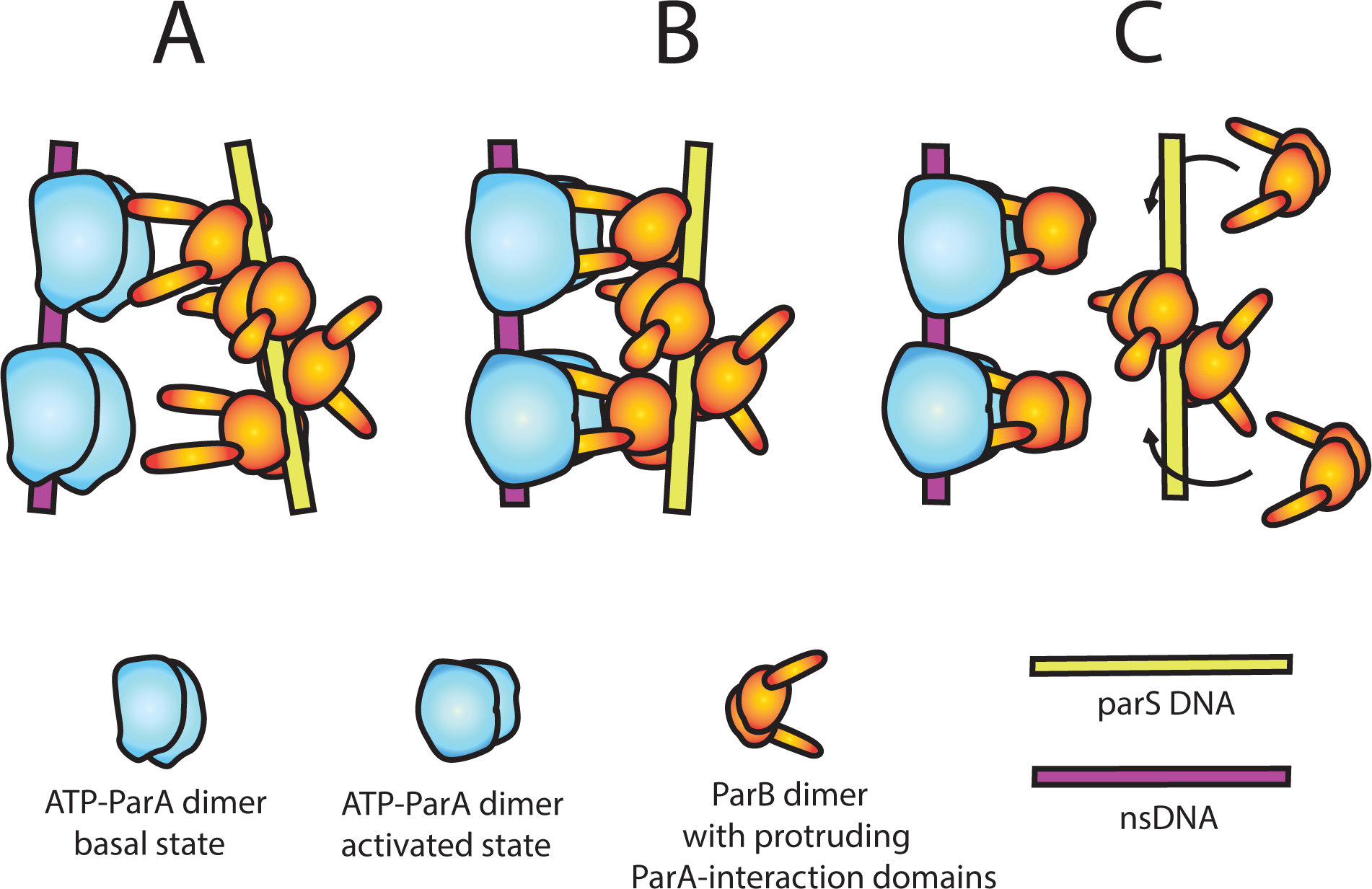
A model of nsDNA-bound ParA_pSM_-ATPase activation by *parS_pSM_*-bound ParB_pSM_. (A) ParA_pSM_-ATP dimers bound to nsDNA in their basal state can interact with ParA-activation domains protruding from ParB_pSM_ dimers bound to *parS_pSM_* sequence repeats. However, multiple ParA_pSM_ dimers in a mini-filament cannot simultaneously interact with ParB_pSM_ dimers bound to a set of repeated sequence copies within a *parS_pSM_* site and the two proteins dissociate quickly. We propose the interacting pair of proteins at this stage are not fully engaged and these ParA_pSM_ dimers are not in the ATPase-activated state. Here, we assume two ParA-interacting domains belonging to one ParB_pSM_ dimer binds one ParA_pSM_ dimer. (B) We propose torsional (or other conformational) thermal Brownian dynamics of the mini-filaments allow the ParB_pSM_ dimer at the fourth position to establishes interaction with another ParA_pSM_ dimer, locking in the non-equilibrium conformation of the individual mini-filaments prior to the formation of this second bridge. The distortion promotes conformational transition of the ParA_pSM_ dimers to the ATPase active state with fully engaged ParB_pSM_. (C) The conformational transition also destabilizes ParB_pSM_-*parS* interaction, releasing the *parS_pSM_* DNA from the activated nsDNA-ParA_pSM_-ParB_pSM_ complex prior to ATP hydrolysis and disassembly of the complex. This allows reloading of fresh ParB_pSM_ to the released *parS_pSM_*, which recycles to disassemble the remaining ParA_pSM_ on the nsDNA. Meanwhile, fully engaged ParB_pSM_ dimers left on the ParA_pSM_ dimers cause ATP hydrolysis and disassemble ParA_pSM_ dimers from nsDNA.

The concentration mismatch between ParB_pSM_ and *parS_pSM_* needed for full activation of ParA_pSM_ ATPase (Fig. 1D) suggests additional details; the functional *parS_pSM_* DNA might be needed only to deliver ParB_pSM_ dimers onto ParA_pSM_-nsDNA mini-filament generating a complex committed to ATP hydrolysis. After ParB_pSM_ delivery, *parS_pSM_* might release the delivered ParB_pSM_ dimers, perhaps helped by the mini-filament distortion discussed above, without waiting for ATP hydrolysis (Fig. 6C). The emptied consensus sequence on the *parS_pSM_* would then be quickly recharged with free ParB_pSM_ dimers in solution to activate another ParA_pSM_-nsDNA complex, explaining the higher concentration demand for ParB_pSM_ over *parS_pSM_* for ATPase activation. The requirement for *parS_pSM_* and ParB_pSM_ replenishment from the washing solution to sustain maximum *k_off_* of the nsDNA-bound ParA_pSM_ dimers (Fig. 4C) supports the notion that the initial active ParA_pSM_-ParB_pSM_-*parS_pSM_* complex assembled on the nsDNA perhaps does not induce ATP hydrolysis of all the ParA_pSM_ dimers within the complex. Full disassembly of the complex likely involves release of the *parS_pSM_* fragment, which gets reloaded with ParB_pSM_ in solution and returns to the ParA_pSM_ dimers left on the nsDNA. This local recycling of ParB_pSM_ and *parS_pSM_* would become inefficient when ParB_pSM_ and *parS_pSM_* are removed from reaction by the buffer wash. We note that dependence of the accelerated phase of ParA_pSM_ *k_off_* on the concentration of the *parS_pSM_*- ParB_pSM_ complex (Table 1) is consistent with the above observation. *In vivo*, local off-rate of ParA_pSM_ from nucleoid where a PC is located is likely to be significantly faster than observed here, considering the local density of the nucleoid-bound ParA_pSM_ near a PC would be lower and the ParB_pSM_-*parS_pSM_* concentration of a PC, with 6 copies of *parS* sites, much higher compared to the conditions in this study.

Above, we described our model considering one ParB_pSM_-*parS_pSM_* mini-filament acting on one ParA_pSM_ cluster on nsDNA for simplicity. However, this does not immediately explain the clearly biphasic disassembly kinetics of the nsDNA-bound ParA_pSM_ during washing with ParB_pSM_-*parS_pSM_* complex (Fig. 2B, 3Aa, e, i). This kinetics is highly reminiscent of activation and membrane dissociation kinetics of membrane-bound MinD-ATPase, a member of ParA-ATPase family, when washed by MinE-containing buffer (Vecchiarelli et al., 2016), which has been proposed to reflect requirement for two MinE dimers for the activation of one MinD dimer. Thus, we suspect two ParB_pSM_-*parS_pSM_* mini-filaments may need to cooperate from two sides of one ParA_pSM_ cluster on nsDNA, and the second binding is kinetically limiting.

This study revealed a puzzlingly unique *parS_pSM_* DNA site structure required for the assembly of the fully active ParB_pSM_-*parS_pSM_* PC in the pSM19035 ParABS system. We propose one possible scenario to explain our findings. However, the model presented here is perhaps not the only possible explanation. Further studies are needed to support or refute the model to advance our mechanistic understanding of this family of partition systems. Further development of multi- faceted approaches to study the synthetic dynamics of the reaction components is needed to advance our mechanistic understanding of the rich variations of the prokaryotic chromosome/plasmid partition systems.

## Materials and methods

### Materials availability

Newly generated materials from this study are available by request to the corresponding author, Kiyoshi Mizuuchi until the lab stocks become exhausted or the lab group operation becomes terminated.

### Proteins peptides and DNA

Non-fluorescent ParA_pSM_-His_6_ (wild-type and mutants) was purified as previously described (Pratto et al. 2008). ParA_pSM_-GFP-His_6_ and ParA_pSMD60E_-GFP-His_6_ were purified as described for ParA_F_-GFP-His_6_ (Vecchiarelli et al. 2010). ParB_pSM_ wild-type and ParB_pSM_-cys, which had three residues (-GCE) added at the C-terminal were purified essentially as previously described with the addition of reducing agent (2mM DTT) in the buffers 50 mM Tris-HCl pH 7.5, 100 mM NaCl plus 5% glycerol. Protein concentrations were estimated based on OD_280_ and aromatic amino acid content.

Fluorescence-labeling of ParB_pSM_ was done as described for Alexa647-ParB_F_ (Vecchiarelli et al. 2013). ParB_pSM_-GCE, in 50 mM Tris-HCl pH 7.4, 100 mM NaCl, 0.1 mM EDTA, 10% glycerol, was mixed with Alexa647-maleimide (Invitrogen) at a protein to dye ratio of 1:1 and then incubated for 1 h at room temperature in the dark. 20 mM DTT was added to stop the reaction. Free dye was removed by spin gel-filtration in a G-50 column. The dye labeling efficiency was determined by spectrophotometry to be ∼15%, and the labeled protein was mixed with unlabeled protein to prepare 1%-labeled ParB_pSM_-Alexa647used for the experiments. *In vivo*, ParB_pSM_-GCE in combination with ParA_pSM_-GFP was fully competent for partition (Maria Moreno-del Álamo, personal communication), and *in vitro*, dye-labeling did not affect its activities, as measured by its ability to stimulate ParA_pSM_ ATPase and by its *parS_pSM_* DNA binding activity (A. Volante, unpublished results).

The N-terminal peptides of ParB_pSM_ (residues 1-27) and ParB_P1_ (residues 1-30) used in the ATPase assays were synthesized by GenScript. The sequence of ParB_pSM1-27_ and its variant ParB_pSM1-27 K10A_ were NH_2_-MIVGNLGAQKAKRNDTPISAKKDIMGD-CO_2_H (≥97 % purity) and NH_2_-MIVGNLGAQAAKRNDTPISAKKDIMGD-CO_2_H (≥ 96 % purity) respectively. The sequence of ParB _1-30_ was NH_2_-MSKKNRPTIGRTLNPSILSGFDSSSASGDR-CO_2_H (≥ 97 % purity).

Two strands of each double-stranded DNA containing heptad repeat (5’-WATCACW-3’) or non-consensus scrambled sequences were synthetized, annealed and purified by ITD (Integrated DNA Technologies). The forward sequences of a DNA duplex are listed in the table below:

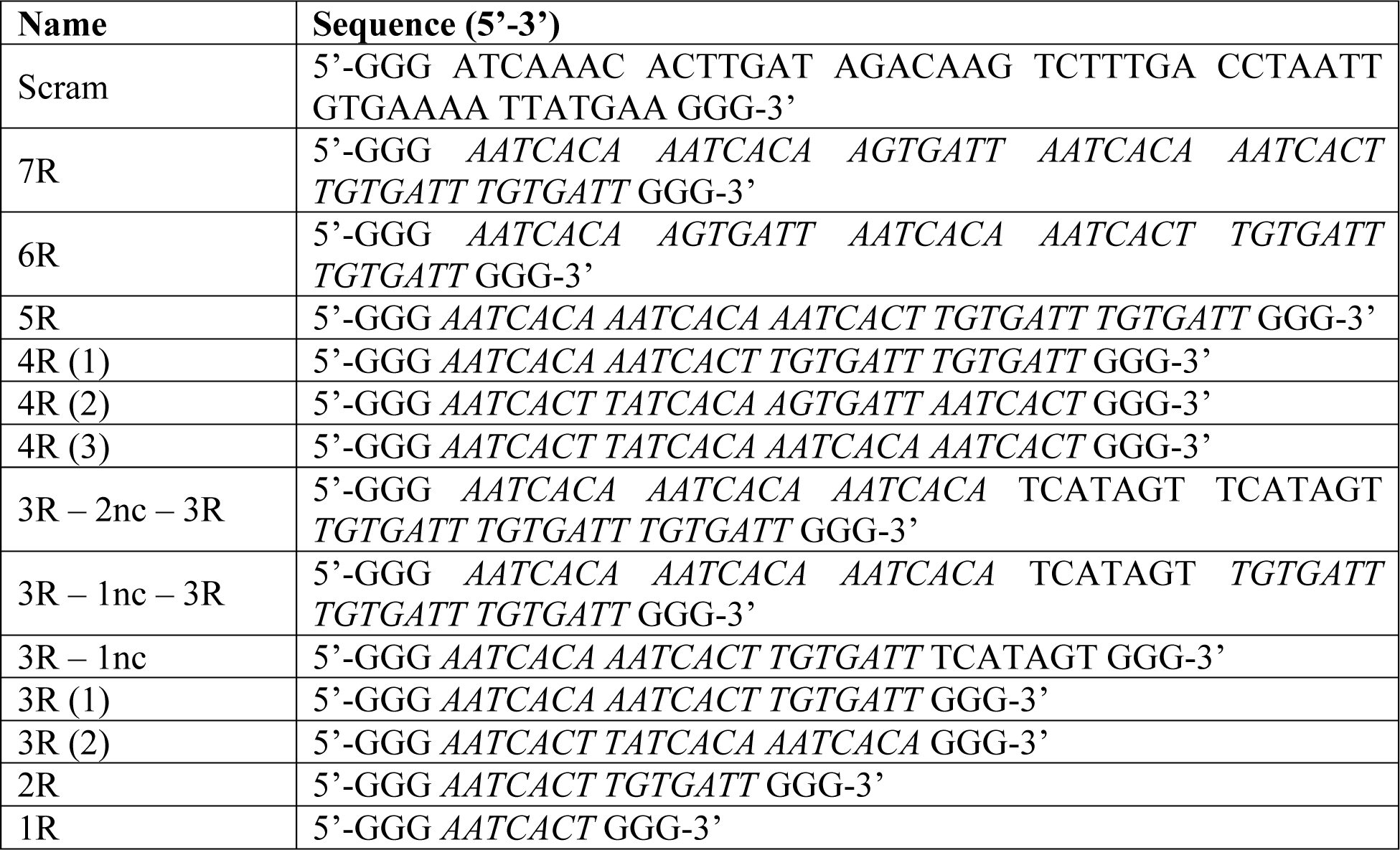

### Plasmids

Plasmids encoding wild-type ParA_pSM_ (Delta, pCB746), mutant ParA_pSMD60A_ (pCB755) and ParB_pSM_ (Omega, pT712ω) have been described previously (Welfe et al. 2005, Pratto et al. 2008, Volante and Alonso, 2015). The pET11a harboring the sequence of his-ParA_pSMD60E_ (pET11a-ParA_pSMD60E_) his-ParA_pSM_-eGFP (pET11a-ParA_pSM_-eGFP) and its variants (D60A, pET11a- ParA_pSMD60A_-eGFP and D60E, pET11a-ParA_pSMD60E_-eGFP) were constructed by GenScript. The ParB_pSM_-GCE [72G, 73C, 74E] allele was created by site-directed mutagenesis and then cloned into the ParB_pSM_ expression plasmid (pT712) to generate pCB1033.

### ATPase assays

Unless stated otherwise, all ATPase assays were performed as follows: the reaction contained 2 µM ParA_pSM_, the indicated concentration of full-length ParB_pSM_ (or ParB_pSM1-27_), nsDNA (plasmid pBR322 DNA, 60 µM in bp) and *parS_pSM_* DNA duplex in 50 mM Tris-HCl (pH7.4), 100 mM KCl, 2 mM MgCl_2_, and 1.5 mM [γ-_32_P]ATP. Labeled ATP was purified before use as previously described (Vecchiarelli et al. 2010). Reactions were incubated at 37°C for 3 h and analyzed by TLC as previously described (Pratto et al. 2008). The data points shown are the means with error bars (standard errors of mean) of repeat experiments (N) as indicated for each figure panel. N* indicates repeat number for majority of ParB_pSM_ concentration points as detailed in the Source Data File. Data sets of repeated measurements were fit after subtraction of background measured without ParA_pSM_ to a modified Hill equation: v – v_0_ = (v_max_ [B]_n_)/(K_An_ + [B]_n_), and the fit parameters and symmetrized error ranges reflecting the larger error of the 95% confidence interval below and above the mean were estimated using Prism 9 (GraphPad). For [B], total concentration of ParB_pSM_ was used instead of free ParB_pSM_ concentration due to technical reasons.

### nsDNA-carpeted flow cell preparation

The flow cells coated with lipid bilayer with attached biotin were prepared essentially as described in (Han and Mizuuchi 2010), and rinsed with a buffer containing 25 mM Tris-HCl pH7.4, 150 mM NaCl and 5 mM MgCl_2_ and 0.1 mM CaCl_2_. Sonicated and biotinylated DNA was prepared as follows: 250 μl of 10 mg/ml sonicated salmon sperm DNA (Sigma) was sonicated for an additional 5 min (Misonix sonicator 3000, output level 6, pulsed on/off for 10 s each at 16°C) to size-weighed average length of ∼500-bp. In order to biotinylate the DNA ends, the sonicated DNA (1 mg/ml) was incubated with 40 μM biotin-17-dCTP (Invitrogen) and 0.6 units TdT (NEB) in the buffer specified by the enzyme manufacturer at 37 °C for 30 min. The reaction was stopped by heating at 70°C for 10 min, and unincorporated biotin-17-dCTP was removed by using S-200 HR Microspin columns (GE Healthcare). The DNA was ethanol precipitated and resuspended in TE buffer. To coat the flow cell with sonicated DNA, the DNA prepared as above was dissolved to 1 mg/ml in 25 mM Tris-HCl pH7.4, 150 mM NaCl, 5 mM MgCl_2,_ and 0.1 mM CaCl_2_, infused into the assembled flow cell, and incubated overnight at 4°C. Unbound DNA was removed by rinsing with 50 mM Tris-HCl (pH 7.4), 100 mM KCl, 2 mM MgCl_2_ and 10% Glycerol.

### TIRFM Setup, Image Processing

Total Internal Reflection Fluorescence (TIRF) illumination and microscopy, as well as the camera settings were essentially as described (Hwang et al. 2013). Combined beam of a 488 nm diode-pumped, solid-state laser (Coherent) and a 633 nm HeNe Laser (Research Electro-Optics) was pointed through a fused silica prism onto the topside of the sample flow cell. Fluorescence emission was collected through a 40× Plan Apo VC, NA 1.4 oil-immersion objectives (Nikon) and magnifier setting at 1.5×. The laser excitation lines were blocked below objective lens with notch filters (NF03-488E and NF03-633E, Semrock). The fluorescence images were captured by an EMCCD camera (Andor IXON+ 897) through a Dual-View module (630DCXR cube, Photometrics; short/long pass filters, SP01-633RS, LP02-633RE, Semrock). Microscopy experiments were carried out at room temperature (∼23°C). Typical camera settings were digitization 16–bit at 1 MHz, preamplifier gain 5.2, vertical shift speed 2 MHz, vertical clock range: normal, EM gain 40, EMCCD temperature set at -98°C, baseline clamp ON. Images were acquired at exposure time 100 ms with frame rate 1, 0.4 or 0.2 Hz using Metamorph 7 software (Molecular Devices) and analyzed using ImageJ software (NIH) as described (Hwang et al. 2013). Data were analyzed with Prism 9 (GraphPad).

### Estimation of the nsDNA-carpet-bound protein densities from the observed fluorescence intensities

ParA_pSM_ and ParB_pSM_ density on the nsDNA-carpet (dimers/μm_2_) were calculated from the fluorescence intensity of acquired images essentially according to the procedure described in the Figure S4 legend in Vecchiarelli et al., 2016. The labeled protein samples used in the experiments were diluted to different concentrations (0 to 8 μM) in a buffer (50 mM Tris-HCl pH 7.4, 100 mM KCl, 2 mM MgCl2, 10% Glycerol, 1 mM DTT, 1 mg/ml α-casein, and 0.6 mg/ml ascorbic acid) and infused into flow cell coated with 1,2-dioleoyl-sn-glycero-3-phosphocholine. Fluorescence intensity data were collected before the protein arrival (background), with the protein sample in the flow cell, and after washing the flow cell with buffer (background) using the illumination and image acquisition parameter settings used for the series of experiments for which the conversion parameters were prepared. The fluorescence signal from the protein sample in the solution was obtained by subtracting the background signal with buffer only (mostly camera dark noise). From the wavelength, the refractive indices of fused silica slide glass and the reaction buffer, and the illumination angle, the evanescence penetration depths were calculated to be 131 and 170 nm for 488- and 633-nm excitation beams, respectively. The protein concentration and the evanescence penetration volume yielded the number of protein molecules, when bound to the flow cell surface, would produce the observed fluorescence signal, yielding the conversion factor used.

### ParA_pSM_ and ParB_pSM_ association and dissociation from the nsDNA-carpet

Two inlet flow cells were assembled and coated with sonicated salmon sperm DNA as described (Vecchiarelli et al. 2013). One inlet was connected to syringe-A containing “binding solution” and the other to syringe-B containing “wash solution”. For protein binding to the nsDNA-carpet, syringe-A delivered at 5 μl/min, while syringe-B at 0.5 μl/min. To start the wash, the flow rates of the two syringes were reversed keeping the total flow rate constant. Experiments were done at room temperature in pSM19035 buffer: 50 mM Tris-HCl (pH 7.4), 100 mM KCl, 2 mM MgCl_2_, 10% Glycerol, 1 mM DTT, 0.1 mg/ml α-casein, 0.6 mg/ml ascorbic acid and 1 mM ATP. Additional components of the binding and washing solutions were as specified in the figure legends. Syringe components were preincubated before infusion at 23°C for 30 min or longer. The association and dissociation data points represent the means with error bars (standard errors of mean) of repeat experiments (N) as indicated for each panel. The dissociation curves were fitted to a single- or a double- exponential equation after subtraction of background in the absence of protein with plateau level set to zero when necessary to avoid large plateau values, and the fraction of fast dissociating population, the *k_off_* and symmetrized error ranges reflecting the larger error of the 95% confidence interval below and above the mean were estimated using Prism 9 (GraphPad).

## Acknowledgements

We are grateful to our colleagues Keir Neuman, Barbara Funnell, Anthony Vecchiarelli, James Taylor, Michiyo Mizuuchi, Min Li, Masaki Osawa and William Carlquist for helpful suggestions and discussion, and to Maria Moreno-del Álamo for providing the information concerning the *in vivo* functionality of ParA_pSM_-GFP/ParB_pSM_-GCE. This work was supported by the intramural research fund for National Institute of Diabetes and Digestive and Kidney Diseases (K.M.), and by the Ministerio de Ciencia e Innovación, Agencia Estatal de Investigación (MCIN/AEI)/FEDER, 2018-097054-B-I00 and 2021AEP031 to JCA.

## Figure Supplements and Legends

**Figure 1-figure supplement 1.**
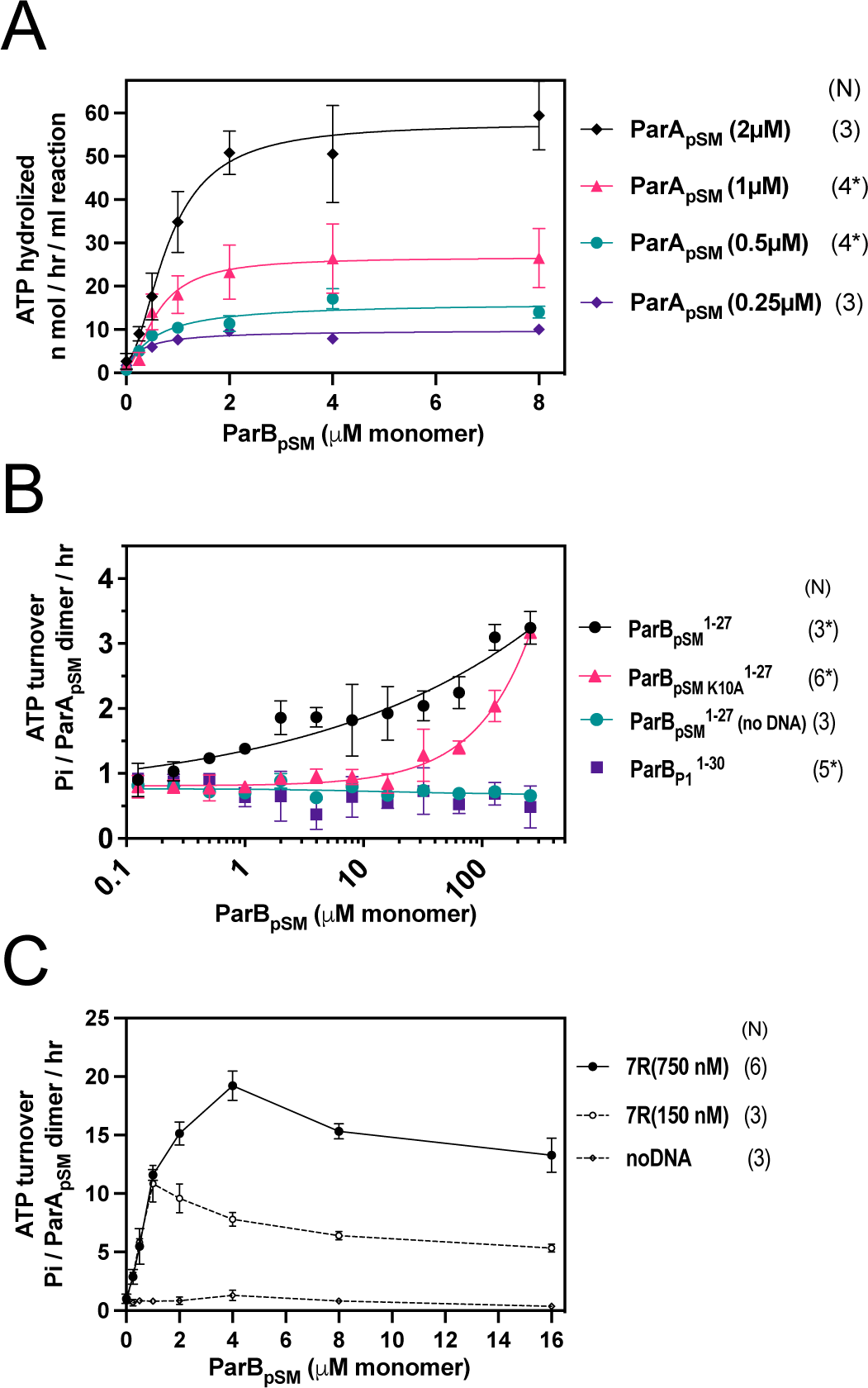
ParA_pSM_-ATPase stimulation: by ParB_pSM_ at different ParA_pSM_ concentrations, by ParB_pSM1-27_, and by ParB_pSM_-7R-*parS_pSM_* in the absence of nsDNA. (A) The ATPase reactions contained ParA_pSM_ at concentrations indicated, nonspecific plasmid DNA (30μM in bp), ParB_pSM_ (at the concentration indicated) and 7R-*parS_pSM_* duplex (34.6 μM in bp or 4.4 μM of the ParB_pSM_-binding consensus sequence). (B) The ParB_pSM1-27_ (black circles) and its variant K10A (red triangles) weakly stimulated ParA_pSM_ ATPase (2μM) in the presence of nsDNA. No stimulation was observed with ParB _1- 30_, which specifically interacts P1 plasmid ParA_P1_ (purple square) or when nsDNA was omitted from the reaction (green circle). (C) Reactions with two concentrations of 7R-*parS_pSM_* dsDNA fragment, 0.75 μM (filled circles) and 0.15 μM (open circles) in *parS_pSM_* fragment concentration (or ∼5 μμ and ∼1 μμ of the ParB_pSM_-binding consensus sequence) in the absence of nsDNA were compared in the presence of a range of ParB_pSM_ concentrations indicated. Data points represent means and standard errors of mean (SEM) of N repeated experiments (N* represents repeats for majority of data points, see Source Data File for details.) Figure 1-Source Data 1

**Figure 1-figure supplement 2.**
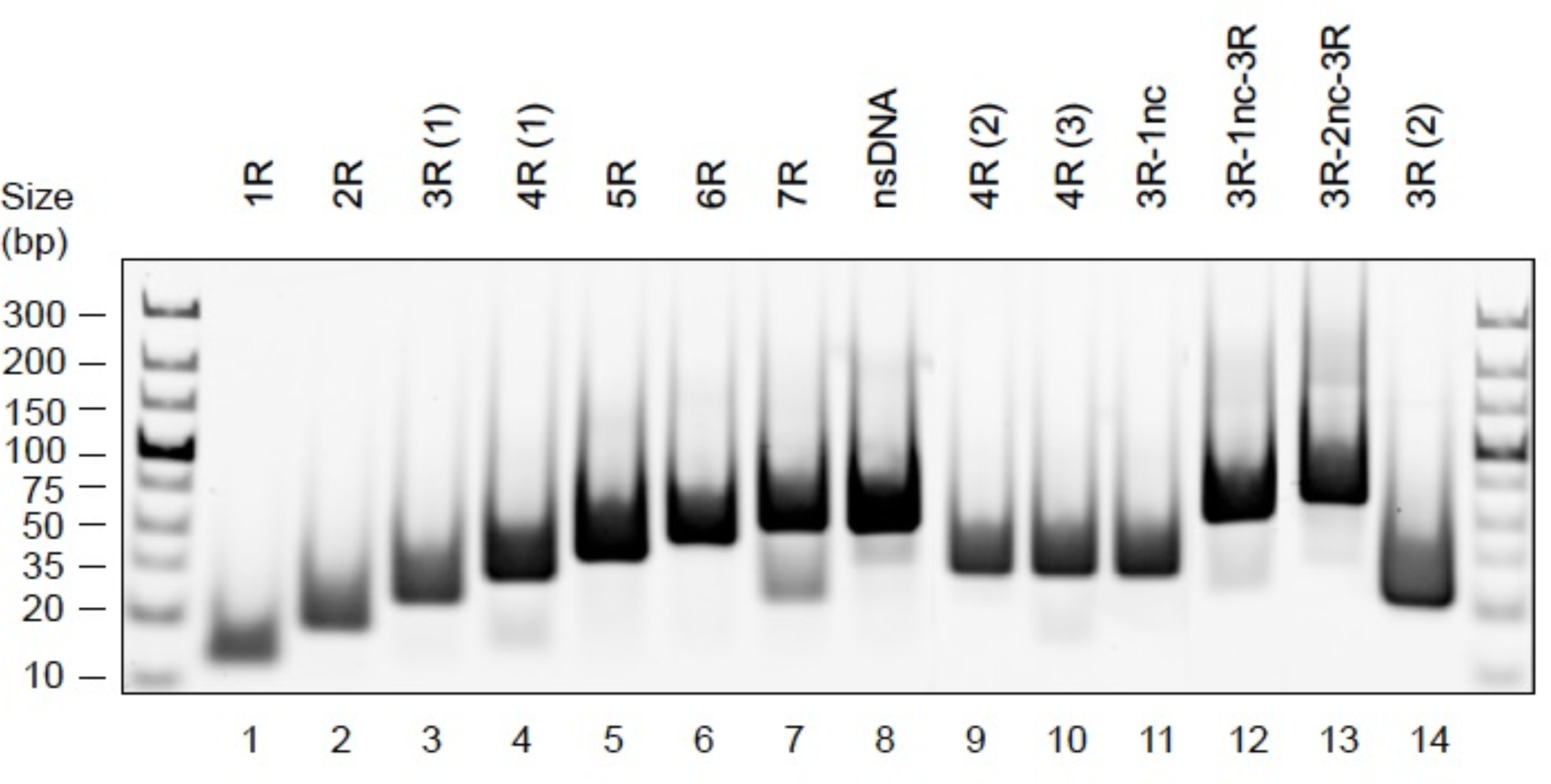
DNA duplex substrates. Analysis of annealed dsDNA fragments (500 ng per well) used in this study by 6% PAGE stained with EtBr.

**Figure 1-figure supplement 3.**
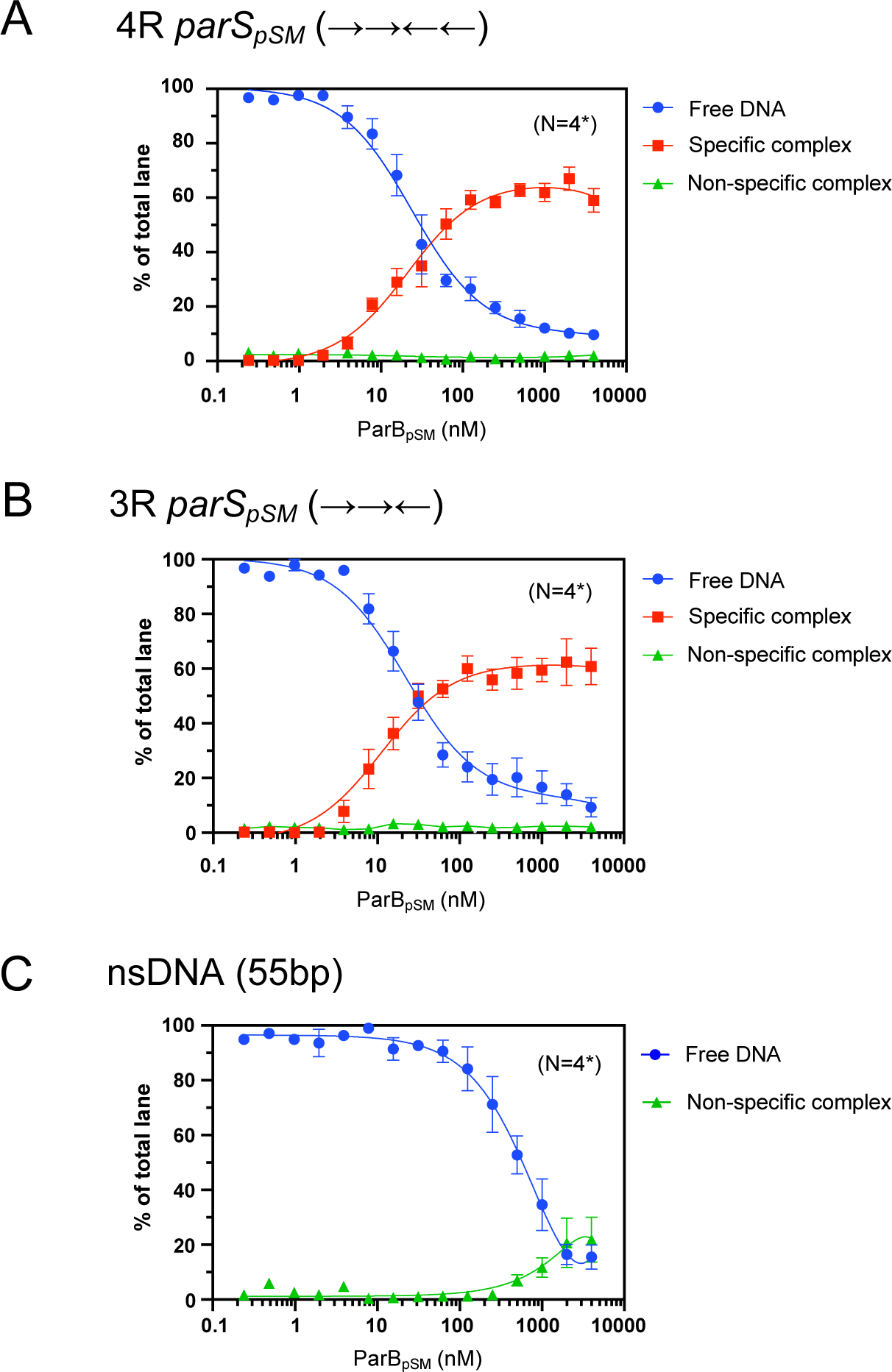
Binding affinity of ParB_pSM_ to 3R- and 4R-*parS_pSM_*. Binding affinity of ParB_pSM_ to 3R- and 4R-*parS_pSM_* was compared to a scrambled sequence DNA fragment (55 bp, nsDNA) by EMSA. Indicated concentrations of ParB_pSM_ was preincubated with 0.05 nM _32_P-labeled DNA substrate in 50 mM Tris-HCl pH7.4, 100 mM KCl, 2 mM MgCl_2_, 1 mM DTT at room temperature for 15 min. The samples were analyzed by 4-12 % PAGE and autoradiography. The smear between the specific complex and free DNA bands in the gel due to dissociation of the complex during electrophoresis was not included in the analysis but assumed to be proportional to the measured quantity of the specific complex. Significant quantity of non-specific complex, which stayed at the top of the gel, was detected only at the highest concentrations of ParB_pSM_ with nsDNA. Figure 1-Source Data 1

**Figure 2-figure supplement 1.**
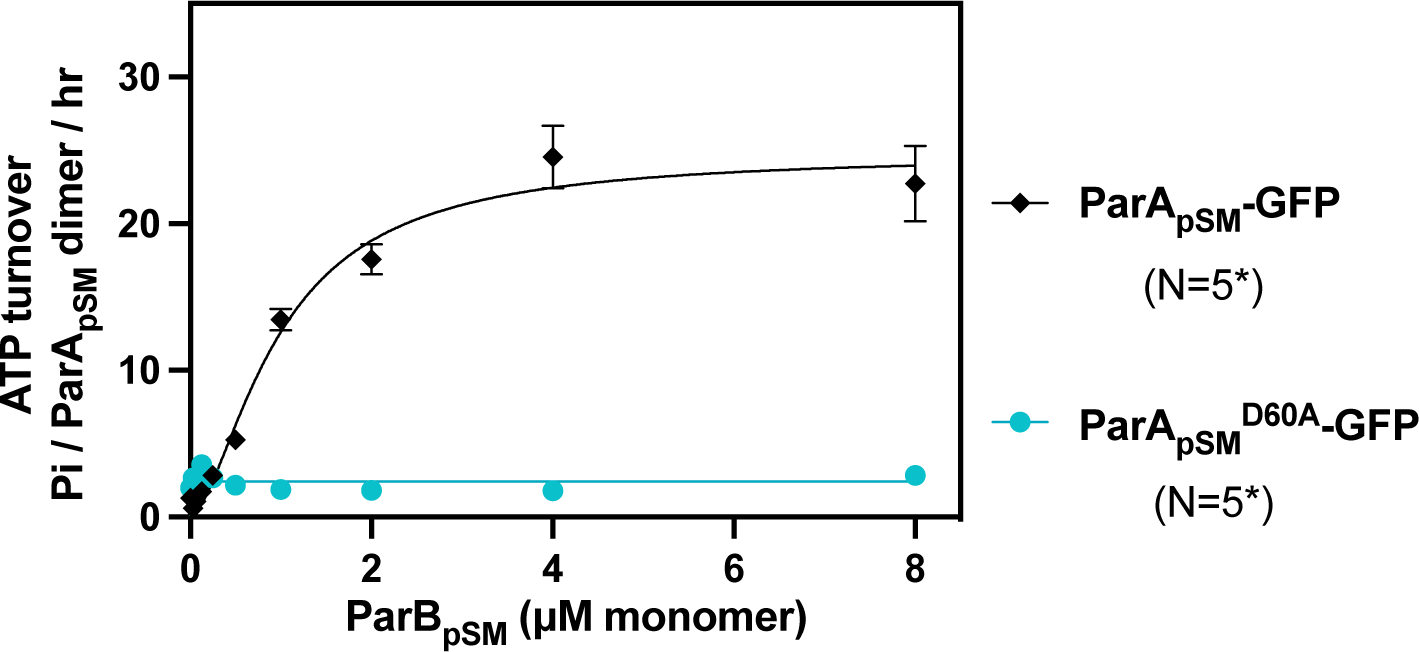
ATPase activity of ParA_pSM_-GFP and the hydrolysis-deficient ParA_pSMD60E_-GFP. The ATPase reaction contained 2 μM ParA_pSM_-GFP (black) or ParA_pSMD60E_-GFP (cyan), pBR322 DNA (30μM bp), ParB_pSM_-Alexa647 (1% labelled) and 7R-*parS_pSM_* DNA (4.4 μM 7-bp consensus sequence repeats). Measurements are averages of at least three independent measurements and the error bars represent the standard errors of mean. N* indicates some lower concentrations of ParB_pSM_ were not included in some of the experimental repeats. Figure 2-Source Data 1

**Figure 2-figure supplement 2.**
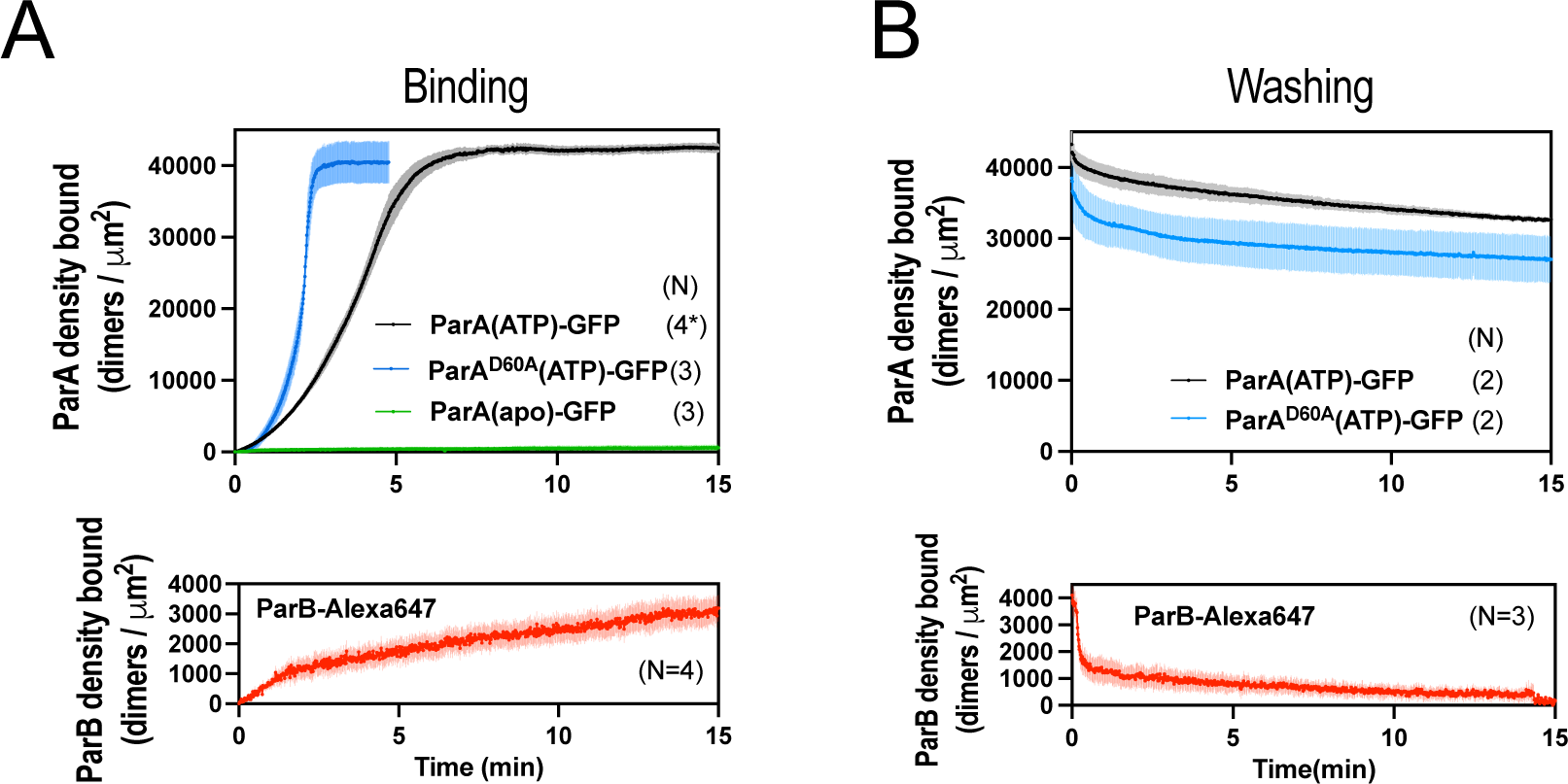
ParA_pSM_, ParA_pSMD60E_ and ParB_pSM_ interactions with the DNA-carpet measured each separately. (A) ParA_pSM_-GFP, ParA_pSMD60E_-GFP (top) and ParB_pSM_- Alexa647 (bottom) binding to the nsDNA-carpet separately. ParA**_pSM_**-GFP (1 μM, black), ParA_pSMD60E_-GFP (1 μM, cyan) or ParB_pSM_-Alexa647 (1% labelled,1 μM, red) were preincubated with 1mM ATP and separately infused into the nsDNA-carpeted flow cell at 5 μl/min for 15 min. ParA_pSM_-GFP (apo) (green line, top) was without ATP. (B) Dissociation kinetics of proteins separately bound to the nsDNA carpet. ParA**_pSM_**-GFP (black) and ParA_pSMD60E_-GFP (cyan) (top) or ParB_pSM_-Alexa647 (bottom, red) densities on the nsDNA-carpet during wash with buffer containing ATP. Each time point represents means and standard errors of mean (vertical spread) of N experiments. Figure 2-Source Data 1

**Figure 2-figure supplement 3.**
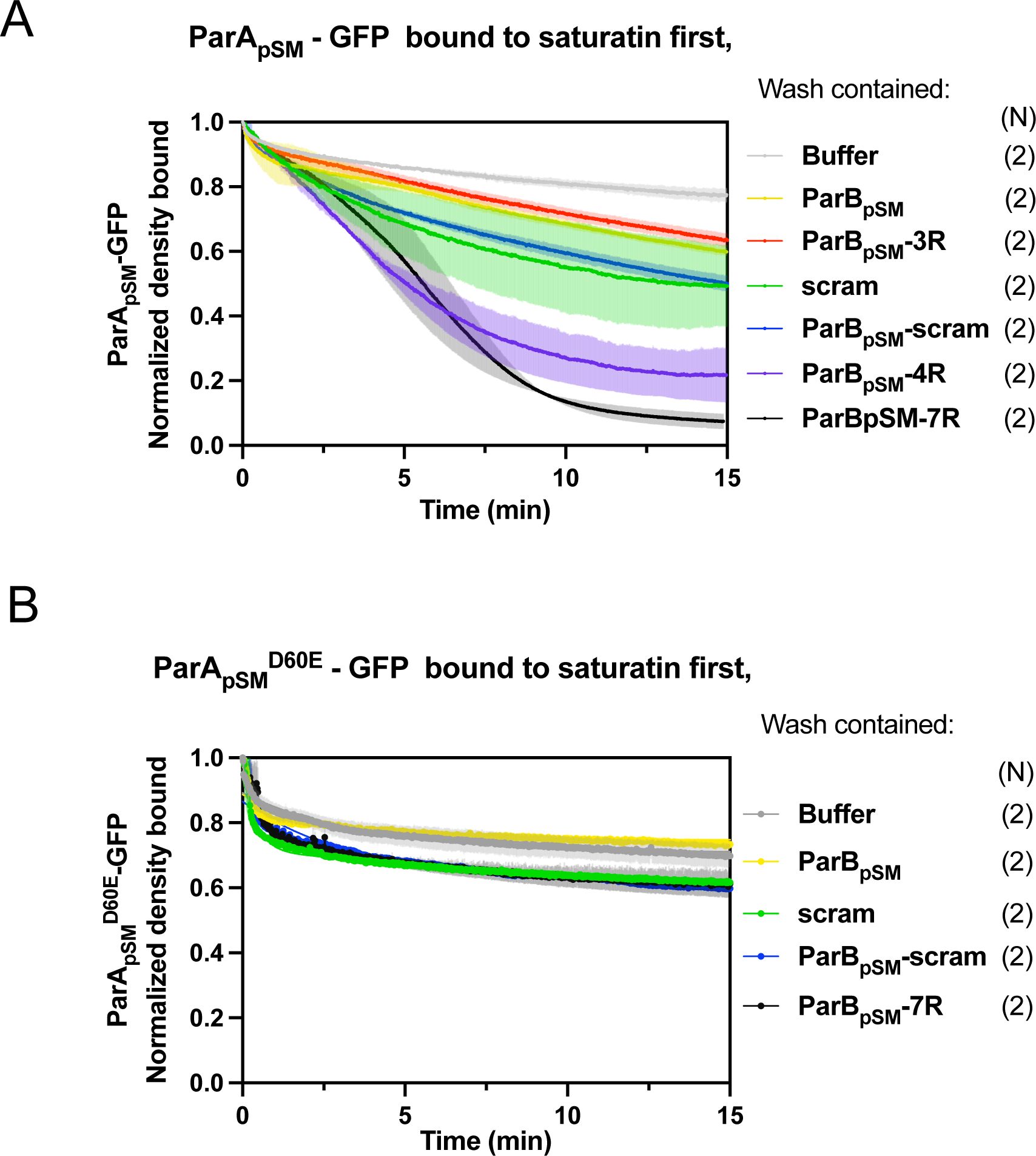
ParB_pSM_-activated dissociation of ParA_pSM_-GFP or ParA_pSMD60E_-GFP bound to nsDNA-carpet to saturation density. ParA**_pSM_**-GFP (A) or ParA_pSMD60E_-GFP (B) (1 μM) was preincubated with 1mM ATP and infused into the nsDNA-carpeted flow cell at 5 μl/min up to saturation binding. At t = 0, the solution flowing over the observation area was switched to the ATP-containing wash buffer with the indicated cofactors: ParB_pSM_, 1 μM with or without; 3R-, 4R-, or 7R- *parS_pSM_* (0.5 μM consensus sequence), or 55-bp scrambled sequence duplex (5.5 μM in bp) as indicated. ParB_pSM_ and *parS_pSM_* fragments were pre-incubated prior to use. The fluorescence intensity of ParA-GFP proteins were normalized to that at t = 0. Each time point represents means and standard errors of mean (vertical spread) of N experiments. Figure 2-Source Data 1

